# Phosphorylation of PLPPR3 membrane proteins as signaling integrator at neuronal synapses

**DOI:** 10.1101/2024.03.11.584206

**Authors:** Cristina Kroon, Shannon Bareesel, Marieluise Kirchner, Niclas Gimber, Dimitra Ranti, Annika Brosig, Kathrin Textoris-Taube, Timothy A. Zolnik, Philipp Mertins, Jan Schmoranzer, George Leondaritis, Britta J. Eickholt

## Abstract

Phospholipid-phosphatase related protein 3 (PLPPR3, previously known as Plasticity Related Gene 2 or PRG2) belongs to a family of transmembrane proteins, highly expressed in neuronal development, which regulate critical growth processes in neurons. Prior work established crucial functions of PLPPR3 in axon guidance, filopodia formation and axon branching. However, little is known regarding the signaling events regulating PLPPR3 function. We identify here 26 high-confidence phosphorylation sites in the intracellular domain of PLPPR3 using mass spectrometry. Biochemical characterization established one of these – S351 – as a *bona fide* phosphorylation site of PKA. Experiments in neuronal cell lines suggest that phosphorylation of S351 does not regulate filopodia formation. Instead, it regulates binding to BASP1, a signaling molecule previously implicated in axonal growth and regeneration. Interestingly, both PLPPR3 intracellular domain and BASP1 enrich in presynapses in primary neurons. We propose that the presynaptic PLPPR3-BASP1 complex may function as novel signaling integrator at neuronal synapses.

## Introduction

Phospholipid phosphatase-related protein 3 (PLPPR3), previously also known as plasticity-related gene 2 (PRG2), belongs to a family of membrane proteins (PLPPR1-5) with described functions in neuronal growth and synaptic transmission.^1^ All PLPPRs share highly conserved six transmembrane domains, and much less conserved intracellular domains of various lengths. This topology makes them ideally suited to integrate cell extrinsic and cell intrinsic signals involved in orchestrating growth and plasticity.

PLPPR3 is highly expressed during early stages of neuronal development, where it controls guidance of thalamic axons and fine-tunes axonal branching by inducing filopodia.^2–4^ In developing glutamatergic neurons, it localizes to the axonal plasma membrane, where it forms clusters, and previous work in our lab has demonstrated that filopodia form at or near these PLPPR3 clusters.^3^ However, our recent data indicate that PLPPR3 expression persists in the adult, where it is highest in the striatum and nucleus accumbens.^5^ Moreover, PLPPR3 is found in synaptosomal fractions isolated from forebrains of adult mice.^5^ Thus, it is likely that PLPPR3 has hitherto unknown synaptic functions.

The functions of PLPPR3 are mediated by its long intracellular domain, which is ideally suited to form signaling complexes. Indeed, PLPPR3-based filopodia formation requires binding and inhibition of the phosphatase PTEN via its intracellular domain, and PLPPR3-dependent thalamocortical targeting of growth cones relies on binding to radixin via its intracellular domain.^2,3^ How these interactions are regulated is currently not known.

Protein phosphorylation is an important regulatory mechanism that essentially controls most cellular processes. It is a rapid and reversible way of regulating proteins post-translationally. The addition of a phosphoryl group, catalyzed by kinases, adds a negative charge to the protein and leads to changes in its binding properties or conformation, thereby leading to a variety of outcomes such as changes in protein localization or turnover.^6–8^ Due to its rapid and pervasive way of changing protein structure and function, phosphorylation is central to intracellular signaling pathways. Curiously, prediction tools suggest that a large number of residues in the PLPPR3 intracellular domain may be phosphorylated (NetPhos3.1; https://services.healthtech.dtu.dk/services/NetPhos-3.1/), indicating an intriguing possibility for PLPPR3 to act as a signaling hub. To date, phosphorylation of PLPPRs has not been studied.

Here, we identify 26 high-confidence phosphorylation sites in the intracellular domain of PLPPR3 using mass spectrometry. We show that PLPPR3 is phosphorylated by PKA at serine 351 and this phosphorylation event regulates its binding to brain acid soluble protein 1 (BASP1, also known as CAP23 or NAP22), a growth-associated signaling molecule. PLPPR3 and BASP1 co-localize in presynaptic clusters along axons and form a synaptic protein complex with hitherto unknown function. We propose PLPPR3 as a novel signaling molecule at neuronal synapses.

## Results

### The intracellular domain of PLPPR3 is highly phosphorylated

PLPPR3 encompasses six transmembrane domains (aa 1-283) and a long intracellular domain (ICD, aa 284-716). The intracellular region of PLPPR3, as well as other PLPPRs, likely mediates the filopodia formation function as well as protein-protein interactions.^1–3,9–12^ However, the regulation of PLPPR3 by post-translational modifications is currently not understood. We tested the presence of phosphorylation, the most pervasive and well-understood post-translational modification, in primary cortical neurons at DIV9, when PLPPR3 expression peaks in culture.^3^ We used Lambda phosphatase to non-selectively remove all phospho-groups in the sample and assessed phosphorylation status by band shift, as phosphorylated proteins typically migrate slower than the non-phosphorylated proteins in SDS-PAGE. These experiments identified a noticeable band shift in the migration of PLPPR3 in response to the phosphatase treatment (Figure 1A), suggesting that PLPPR3 is a phosphoprotein.

**Figure 1.**
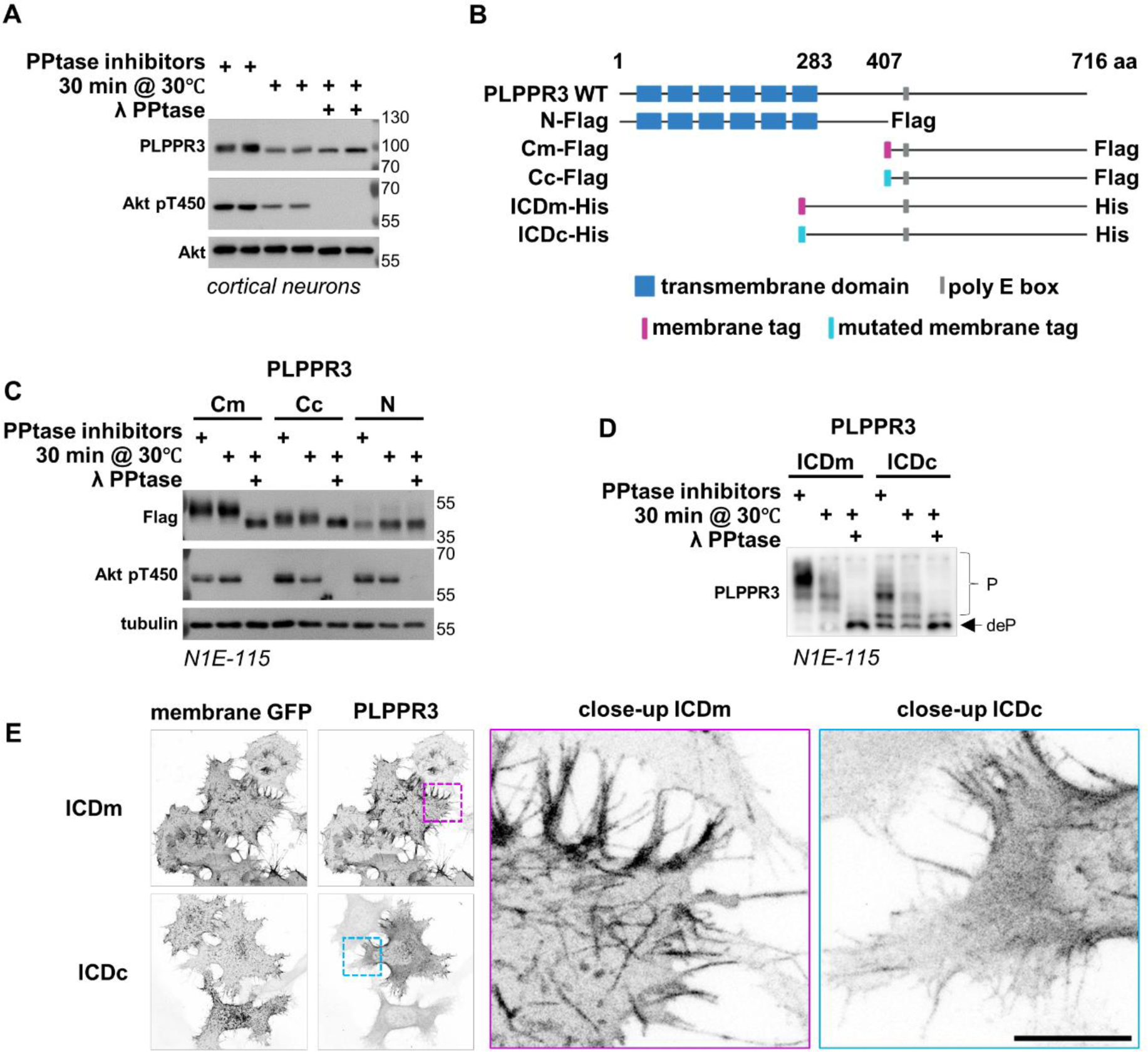
PLPPR3 intracellular domain is phosphorylated. (A) Phosphorylation of endogenous PLPPR3 was assessed in DIV9 primary cortical neurons by band shift on SDS-PAGE in response to dephosphorylation with λ phosphatase (PPtase; 30 minutes at 30°C). (B) Graphic illustration of all PLPPR3 variants used in this study. ICD – intracellular domain. Created with Biorender.com. (C) Phosphorylation of truncated PLPPR3 variants was assessed by band shift on SDS-PAGE in response to dephosphorylation with λ phosphatase. N1E-115 cells were lysed 12 hours after transfection with PLPPR3 variants, and the lysate was dephosphorylated as in (A). (D) Phosphorylation of PLPPR3 ICD variants was assessed in N1E-115 cells by PhosTag SDS-PAGE. (E) Localization of PLPPR3 ICD variants to their intended subcellular compartments. N1E-115 cells were fixed 12 hours after transfection and immunolabelled. Membrane-tagged GFP was used as a marker to assess membrane localization. Scale bar = 10 µm.

We next used the phosphorylation prediction tool NetPhos3.1 (https://services.healthtech.dtu.dk/services/NetPhos-3.1/) to investigate which residues of PLPPR3 could be targeted by phosphorylation. Transmembrane domains and extracellular loops were not considered in our analyses due to their inaccessibility to kinases. Using a stringent probability of >0.75, a total of 43 residues were predicted to be phosphorylated, with 39 of them located in the ICD and four located in the intracellular loops (data not shown). Both the membrane-proximal as well as the distal intracellular region were predicted to include a dense cluster of phosphorylated residues, while the regions adjacent to the polyE box – a unique feature of PLPPR3 consisting of 20 glutamic acids – had fewer. To directly test the accuracy of these predictions, we generated multiple PLPPR3 variants. PLPPR3 N-Flag (aa 1-407) and C-Flag (408-716) roughly divide the full-length protein into N- and C-terminal halves, each containing one of the predicted phosphorylation hotspots. We also generated a mutant encompassing the entire intracellular domain, PLPPR3 ICD (aa 284-716). As PLPPR3 is a transmembrane protein, we hypothesized that proximity to the plasma membrane may play an important role in phospho-regulation of the intracellular domain; therefore, we added either a myristylation/palmitoylation tag to anchor the variants to the plasma membrane (Cm or ICDm), or a mutated myristylation/palmitoylation tag that leaves the variants primarily cytosolic (Cc or ICDc).^13^ All PLPPR3 variants are depicted in Figure 1B.

We expressed the PLPPR3 N- and C-terminal variants in the neuroblastoma cell line N1E-115, which show efficient plasma membrane targeting of PLPPR3 constructs, and analyzed their phosphorylation status using SDS-PAGE followed by western blotting. Curiously, the membrane-tagged PLPPR3-Cm showed a robust band shift in response to the phosphatase treatment, while the cytosolic PLPPR3-Cc showed a smaller, yet still evident band shift, indicating importance of the proximity to the plasma membrane in controlling phosphorylation status (Figure 1C). There was no evidence of phosphorylation in the PLPPR3-N variant.

In order to analyze the phosphorylation of the PLPPR3 intracellular domain in more detail, we made use of the membrane-tagged and cytosolic ICD variants and PhosTag SDS-PAGE analyses. PhosTag is a phosphate-binding molecule that traps these groups in the gel and leads to separation of proteins based on their phosphorylation state.^14^ Employing the Lambda phosphatase treatment as previously, we found that the membrane-tagged ICDm exists in an almost entirely phosphorylated state in cells, as evidenced by multiple strong phospho-bands and a very weak non-phospho band under steady-state conditions (Figure 1D, left three lanes). In contrast, the primarily cytosolic ICDc has fewer higher-order phosphorylation bands, as well as an evident non-phospho band (Figure 1D, right three lanes). Immunolabeling in N1E-115 cells confirmed the localization of these variants to their intended subcellular compartments (Figure 1E). Taken together, these results indicate the presence of multiple phosphorylation events in the intracellular domain of PLPPR3 and suggest the importance of proximity to the plasma membrane in the dynamic regulation of phosphorylation status.

We decided to use this as a strategy to uncover phosphorylation sites in the intracellular domain of PLPPR3 and purified PLPPR3 ICDm and ICDc from membrane and cytosolic fractions of HEK293T lysates using affinity chromatography via the His tag (Figure 2A). Analyses by mass spectrometry identified 26 high-confidence phosphorylation sites in the PLPPR3 intracellular domain (Figure 2B). Half of these phospho-sites were unique to the membrane-tagged PLPPR3 ICDm variant, and one to the PLPPR3 ICDc cytosolic intracellular domain, while twelve phospho-sites were found in both PLPPR3 variants irrespective of their subcellular localization. In contrast to the SDS-PAGE analysis of the PLPPR3-N variant, and in agreement with the prediction data, the membrane-proximal intracellular domain contained 15 phosphorylation sites, with a particularly dense phosphorylation cluster around residues 343-400. In the most distal part of the intracellular region, phosphorylation events were densely clustered between amino acid residues 560-575 (Figure 2C). The high number of phosphorylation sites in eukaryotic proteins has led to the speculation that many of those may be unfunctional, and conservation of phosphorylation sites across species has been proposed as a criterion for identification of functional sites.^15–17^ Thus, we analyzed the conservation of these 26 phosphorylation sites across species from zebrafish to human. As shown in Figure 2B, a large majority of these sites were conserved in five or all analyzed species, indicating that these may represent functional phosphorylation events. Altogether, our experiments demonstrate that the intracellular domain of PLPPR3 is highly phosphorylated.

**Figure 2.**
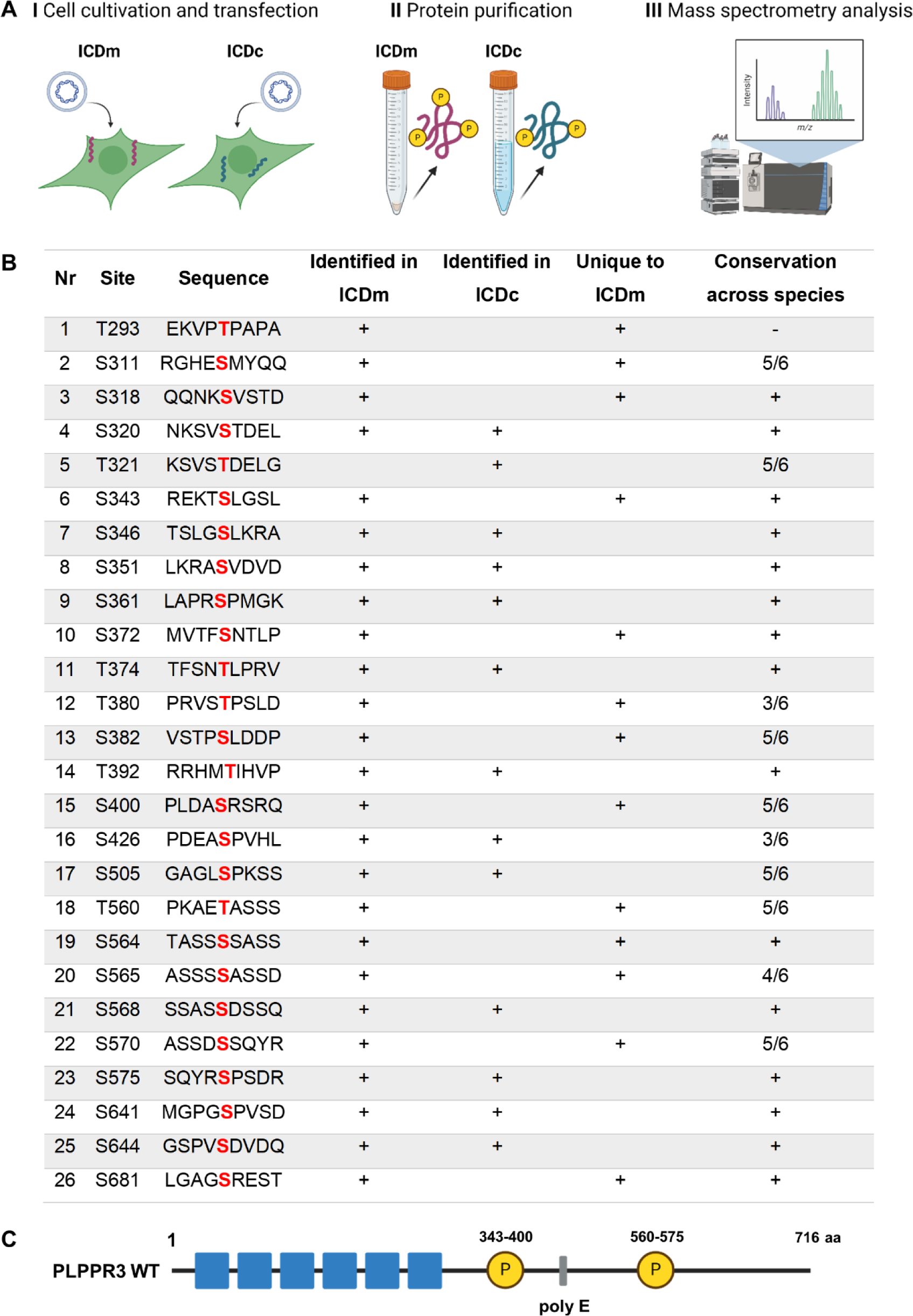
There are 26 high-confidence phosphorylation sites in the intracellular domain of PLPPR3. (A) Experimental workflow. Created with Biorender.com. (B) List of phosphorylated residues in ICDm and ICDc and their conservation across species. Phosphorylated residue is marked in red. Conservation of the phosphorylated residue was analyzed by sequence alignment across six species: mouse (*Mus musculus*), human (*Homo sapiens*), zebrafish (*Danio rerio*), tropical clawed dog (*Xenopus tropicalis*), rhesus monkey (*Macaca mulatta*) and chicken (*Gallus gallus*). Conserved phosphorylation sites are marked with +, nonconserved with -, and for the rest the exact ratio of conserved/total is given. The conservation analysis was performed with COBALT tool (COBALT:Multiple Alignment Tool (nih.gov)). (C) Graphic illustration of PLPPR3 phosphorylation hotspots.

### PLPPR3 S351 is a PKA phosphorylation site

We next investigated which kinases may target the intracellular domain of PLPPR3, focusing on those that (1) had been predicted to target PLPPR3 (NetPhos3.1, data not shown) and (2) that have been described to control filopodia or axon branch formation.^18–21^ We performed an *in vitro* phosphorylation assay using purified PLPPR3 intracellular domain from *E. coli* and commercially available kinases and analyzed protein phosphorylation by PhosTag SDS-PAGE. This analysis identified that protein kinase A (PKA) can phosphorylate purified PLPPR3 intracellular domain, as evidenced by a prominent shift towards higher-order phosphorylation bands (Figure 3A). To test whether PKA can induce PLPPR3 phosphorylation also in cells, we made use of the commercially available PKA modulators depicted in Figure 3B. As a cAMP-dependent kinase, PKA can be activated indirectly by Forskolin, which activates adenylyl cyclases and leads to production of cAMP.^22,23^ Alternatively, PKA can be activated directly by the cAMP analog 8-Br-cAMP, which binds PKA holoenzyme and releases the catalytic PKA subunits.^24^ PKA is inhibited by small molecule inhibitor H89, which competitively binds to the ATP site.^25^ Based on the kinase recognition motif,^26–28^ there are four sites in the PLPPR3 intracellular domain that could potentially be targeted by PKA, and two of them – S351 and T380 – were identified as true phosphorylation sites in our phospho-mass spectrometry analysis (see Figure 2B). However, PLPPR3 S379 has previously been identified as a phosphorylation site in discovery proteomics *in vivo*,^29^ therefore we considered it as well. The fourth site that matches the kinase recognition motif – S581 – was not identified as a phosphorylation site in our mass spectrometry analysis and was therefore not considered.

**Figure 3.**
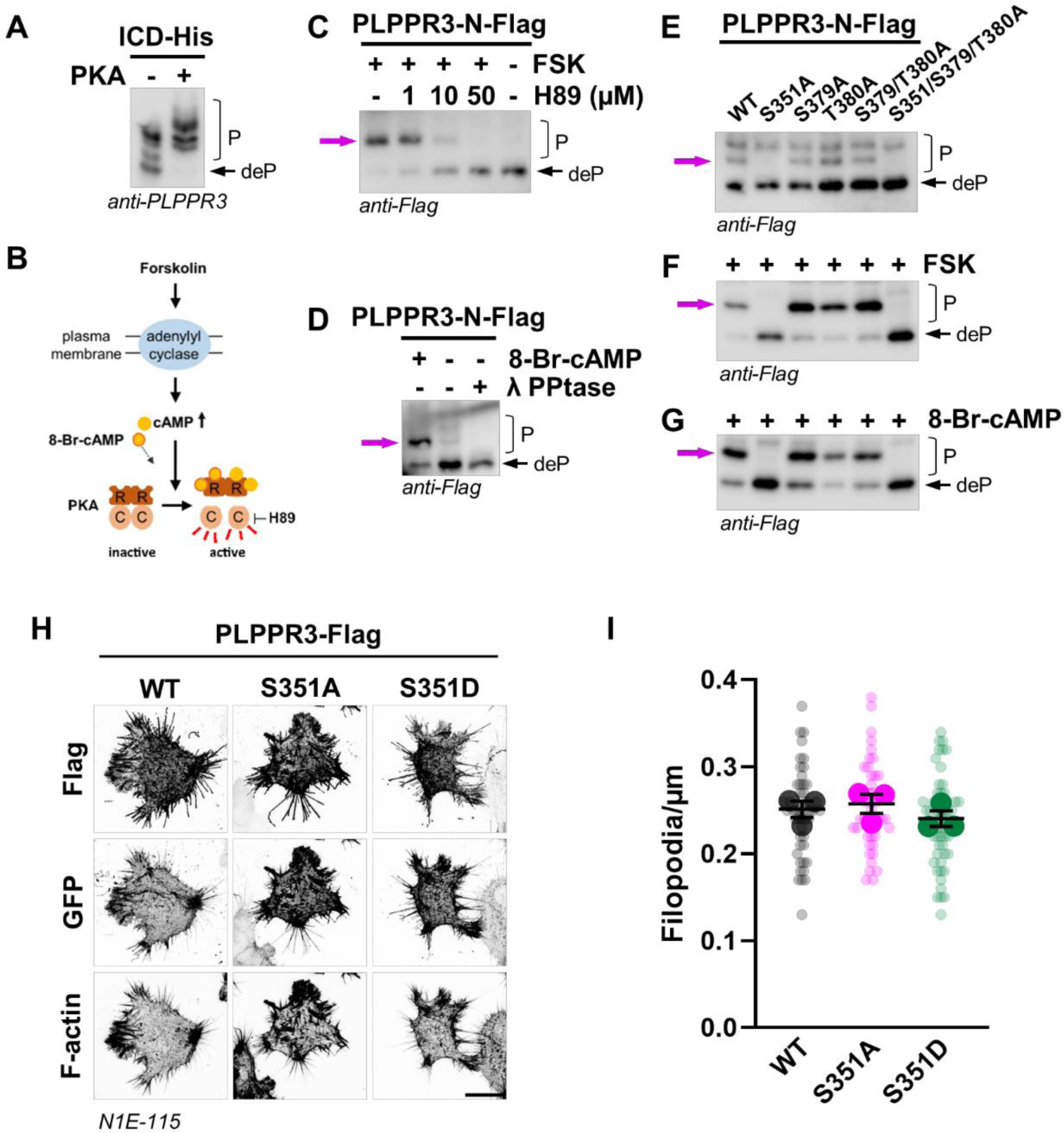
PKA phosphorylates PLPPR3 at S351. (A) In vitro phosphorylation of PLPPR3 ICD by PKA. Purified ICD from E.coli was incubated with PKA in the presence of ATP for 2 hours at 30°C. Samples were analyzed by PhosTag western blot. (B) Graphic illustration of PKA activation in cells and the mechanism of action of drugs used. (C) Phosphorylation of PLPPR3-N-Flag is altered in cells by PKA-modulating drugs. N1E-115 cells were transfected with PLPPR3-N-Flag and serum-starved overnight. Cells were treated with PKA inhibitor H89 for 1 hour, followed by stimulation with 30 µM Forskolin (FSK) for 5 minutes, and analyzed by PhosTag western blot. (D) Phosphorylation of PLPPR3-N-Flag is altered in cells by the second messenger cAMP analog. N1E-115 cells were transfected with PLPPR3-N-Flag and serum-starved overnight. Cells were stimulated with 1 mM 8-Br-cAMP for 5 minutes and analyzed by PhosTag western blot. λ-phosphatase treated sample is included to mark the migration of the unphosphorylated protein. (E) Phosphorylation pattern of PLPPR3-N-Flag phospho-mutants under steady-state conditions. N1E-115 cells were transfected with respective phospho-mutants and protein lysates were analysed by PhosTag western blot. (F, G) Phosphorylation of PLPPR3-N-Flag phospho-dead mutants following activation of PKA. N1E-115 cells were transfected with respective phospho-mutants, stimulated with 30 µM Forskolin (F) or 1 mM 8-Br-cAMP (G) for 5 minutes, and protein lysates were analysed by PhosTag western blot. In all blots, deP indicates migration of the unphosphorylated protein, P phosphorylated forms of protein. Magenta arrows point to the changes in phosphorylation pattern. (H) Filopodia density measurements in N1E-115 cells. Cells were transfected with PLPPR3 phospho-mutants and membrane-tagged GFP, and filopodia density was measured using a semi-automated method. Scale bar = 20 µm. (I) Quantification of (H). N=3, n=43-52, mean ± SEM.

All these three potential PKA sites are found in the membrane-proximal intracellular domain. To test whether any of these sites were targeted by PKA in cells, we made use of the PLPPR3-N-Flag variant (see Figure 1B) and tested it by PhosTag analysis. We observed that stimulation with Forskolin leads to phosphorylation of PLPPR3-N, as indicated by the appearance of a phospho-band, while increasing doses of H89 inhibit the phosphorylation event (Figure 3C). Furthermore, 8-Br-cAMP induces the phosphorylation of PLPPR3-N (Figure 3D), indicating that PKA can phosphorylate PLPPR3 in a cAMP-dependent manner. As in previous experiments, we used Lambda phosphatase treatment as reference for the non-phosphorylated form of PLPPR3-N variant. Since PhosTag analysis cannot always assess whether phospho-bands represent individual phosphorylation events, we used site-directed mutagenesis to generate non-phosphorylatable (S→A) mutants of the three potential PKA motifs, as well as a double and a triple mutant, to identify which site(s) PKA targets. We observed that the S351A, as well as the S351A/S379A/T380A triple mutant lacked a phosphorylation band under steady state conditions (Figure 3E). Furthermore, we showed that the S351A and the triple mutant cannot be phosphorylated in response to Forskolin (Figure 3F) or 8-Br-cAMP (Figure 3G). Taken together these results demonstrate that S351 is a PKA phosphorylation site in the PLPPR3 intracellular domain.

PLPPR3, like other PLPPRs, induces filopodia formation, which prompted us to test whether PLPPR3-induced filopodia formation was regulated by phosphorylation of S351.^1,3,4^ We expressed PLPPR3 wild type (WT) protein, which induces filopodia formation in N1E-115 cells,^30^ and compared it to phospho-dead (S351A) and phospho-mimic (S351D) variants. PLPPR3 phospho-mutants showed equivalent plasma membrane localization with the WT protein, which is considered a requisite of PLPPR-induced filopodia formation.^1^ However, there were no differences in filopodia density between the phospho-mutants and the WT (Figure 3H, I). Thus, we concluded that phosphorylation of PLPPR3 S351 does not regulate filopodia formation.

### Phosphorylation of PLPPR3 S351 regulates its binding to BASP1

Although PLPPR3 S351 does not control filopodia formation, conservation analysis suggests that it may be a functionally relevant site (see Figure 2B). Therefore, we decided to raise an anti-PLPPR3 pS351 phospho-site-specific antibody to facilitate functional analyses. Following affinity purification, we rigorously tested the specificity of this antibody towards the PLPPR3 pS351 epitope using a phospho-dead mutant (PLPPR3 S351A), Lambda phosphatase treatments, as well as *Wt* and *Plppr3^-/-^* neuron lysates. Together, these findings indicate that anti-pS351-PLPPR3 specifically recognizes the serine 351-phosphorylated form of PLPPR3 in western blot applications (Supplementary Figure 1). To verify that this site is phosphorylated in neurons, we first analyzed its temporal pattern in primary hippocampal cultures. Total PLPPR3 protein levels are increased between 5-9 days *in vitro*, coinciding with the time axons undergo branching.^3^ PLPPR3 pS351 was phosphorylated at all tested timepoints, overall following the same abundance pattern as total PLPPR3 with increased levels between 5-9 DIV and decreasing thereafter (Figure 4A). However, the increase in pS351 levels was not as robust as in total PLPPR3 and may thus even be suppressed at 5-9 DIV in relation to total PLPPR3.

We next used Forskolin to investigate if phosphorylation of S351 can be triggered in neurons at these timepoints when PLPPR3 protein levels are elevated. While PKA activation triggers increase of pS351 levels at all tested timepoints, the largest increase was observed at 9 DIV (Figure 4B). We then tested whether PKA-dependent S351 phosphorylation can also be induced in adult brain tissue, where structural and functional integrity of intrinsic synaptic connections is maintained.^31^ Indeed, stimulation of an acute brain slice with Forskolin led to strong increase of PLPPR3 pS351 levels, similar to GluA1 pS845, which is a known PKA target (Figure 4C).^32^ Taken together, these experiments confirm that phosphorylation of PLPPR3 at S351 can be dynamically modified in “*in vivo*-like” setting and further reinforce the idea that this site is a functionally relevant regulatory site.

**Figure 4.**
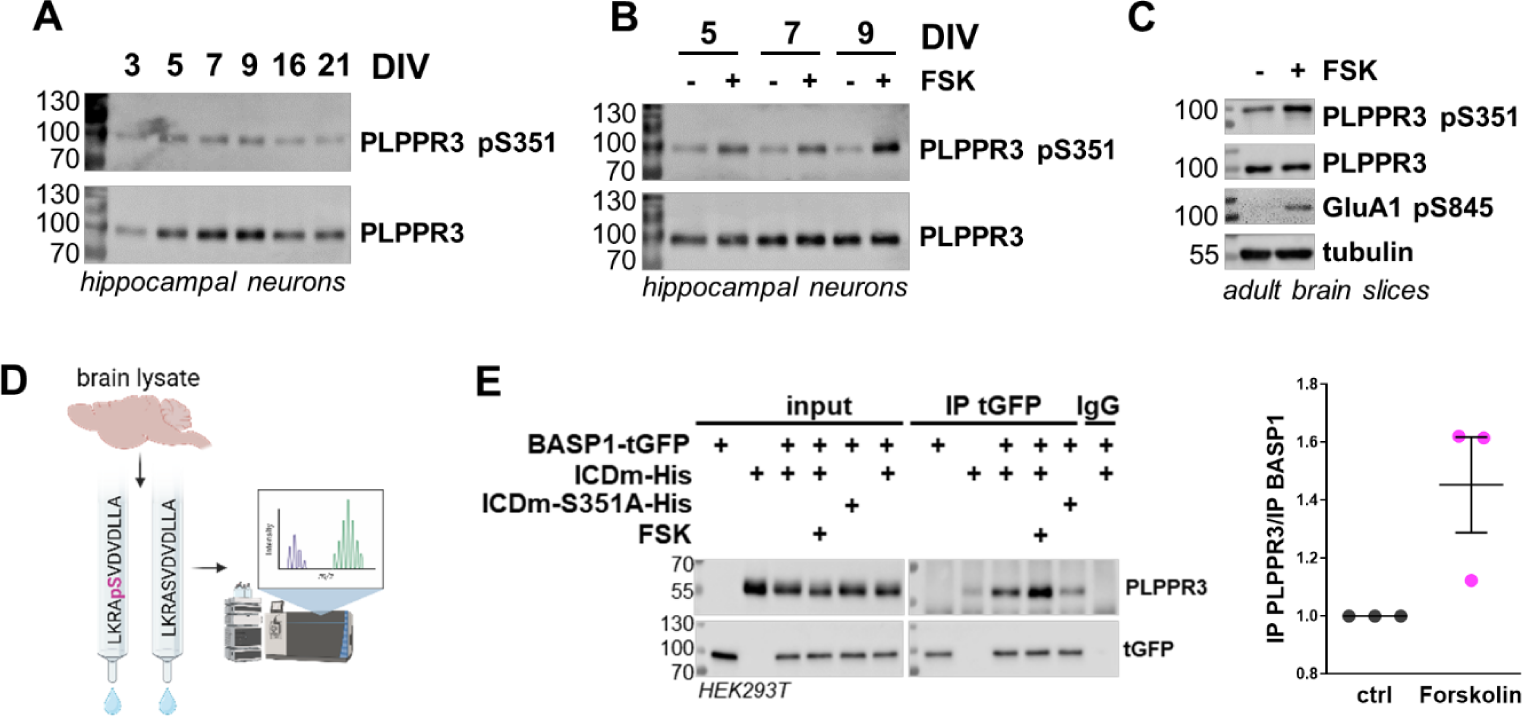
Phosphorylation of PLPPR3 S351 regulates binding to BASP1. (A) PLPPR3 pS351 levels mimic total PLPPR3 levels through developmental stages. Primary hippocampal neurons were lysed at indicated timepoints and pS351 levels were analyzed using custom made antibody. (B) Phosphorylation of PLPPR3 S351 can be triggered in neurons. Primary hippocampal neurons were stimulated with Forskolin (30 µM, 5 minutes) at different timepoints, and analyzed by western blot. (C) Phosphorylation of PLPPR3 S351 can be triggered in adult brain tissue. Acute brain slices were stimulated with Forskolin (30 µM, 15 minutes), lysed and analyzed by western blot. (D) Experimental workflow of the unbiased interaction partner screen. Affinity columns with immobilized pS351 and S351 peptides were incubated with brain lysate 1 hour, eluted, and eluates were analyzed for interaction partners using mass spectrometry. (E) Co-immunoprecipitation of BASP1 and PLPPR3 ICDm. Protein lysates were prepared from HEK293T cells expressing BASP1-tGFP and ICDm-His, and BASP1 was immunoprecipitated using tGFP antibody. Cells were stimulated with 30 µM Forskolin for 10 minutes, where applicable. Quantification of protein bands is seen on the right. The amount of co-immunoprecipitated PLPPR3 ICDm is expressed over precipitated BASP1 amount. All control values were normalized to 1.

Phosphorylation often regulates the nature and strength of protein-protein interactions,^33^ thus we established an unbiased screen to search for interaction partners of PLPPR3 pS351. We used affinity columns conjugated with phospho- (LKRA**pS**VDVDLLA) and non-phospho (LKRA**S**VDVDLLA) peptides to fish out binding partners from adult brain lysate and then characterized the bound proteins using mass spectrometry (Figure 4D). We uncovered three candidate PLPPR3 ICD interacting proteins (Supplementary Figure 2), of which BASP1 closely matched the known spatiotemporal expression as well as the function of PLPPR3.^34–36^ Curiously, BASP1 was robustly enriched in the phospho-column compared to the non-phospho column, suggesting favored binding to phosphorylated S351 peptide. Using co-immunoprecipitation, we directly tested the interaction of PLPPR3 intracellular domain and BASP1. Indeed, PLPPR3 intracellular domain was pulled down with BASP1 under steady state conditions, suggesting binding of the two proteins (Figure 4E). In agreement with increased binding to the phospho-peptide, more PLPPR3 co-immunoprecipitated with BASP1 after Forskolin stimulation. In contrast, the non-phosphorylatable PLPPR3-ICDm-S351A variant showed decreased binding to BASP1. These experiments establish BASP1 as a novel interaction partner of PLPPR3 intracellular domain and indicate that the PLPPR3-BASP1 interaction is regulated by PLPPR3 phosphorylation at S351.

### PLPPR3 and BASP1 co-localize in presynaptic clusters along axons

BASP1 is a growth-associated protein described to regulate axon development and regeneration, and it has been shown to induce filopodia in non-neuronal cells.^34,37^ Thus, we initially investigated the co-localization of PLPPR3 intracellular domain and BASP1 by co-expressing the proteins in neuronal and non-neuronal cells. In HEK293T cells, the two proteins co-localized in the cell peripheral structures and occasionally enriched at the tips of filopodia (Figure 5A). In primary hippocampal neurons, PLPPR3 intracellular domain and BASP1 co-localized in distinct puncta along the axons (Figure 5B). These puncta were reminiscent of immature presynaptic clusters, which prompted us to investigate whether they co-localize with known presynaptic markers. Co-labeling between endogenous VGLUT1, a presynaptic marker, and overexpressed BASP1 identified the co-localization of some BASP1 clusters with VGLUT1 clusters (Figure 5C), however, in dense neuronal cultures it was not always apparent whether the signal was truly coming from the same neurons. To overcome this, we co-expressed BASP1 with another presynaptic marker, Synaptophysin-1. As seen on Figure 5D, BASP1 clusters showed high co-localization with Synaptophysin-1 clusters. Furthermore, also PLPPR3 intracellular domain co-localized with Synaptophysin-1 in distinct puncta (Figure 5E). We quantified the localization of PLPPR3 and Synaptophysin-1 clusters and found that 54% of PLPPR3 clusters are found within presynaptic loci (Figure 5F). Finally, we used crude synaptosomes prepared from adult mice to investigate the localization of endogenous PLPPR3 and PLPPR3 pS351 to synaptic fractions. As seen on Figure 5G, both PLPPR3 as well as PLPPR3 pS351 enriched almost exclusively in synaptosomal fractions, as did known synaptic markers PSD-95 and Synaptophysin-1. Taken together, these experiments characterize PLPPR3-BASP1 as a novel presynaptic protein complex under the control of PKA via direct phosphorylation of PLPPR3 S351 (Figure 5H).

**Figure 5.**
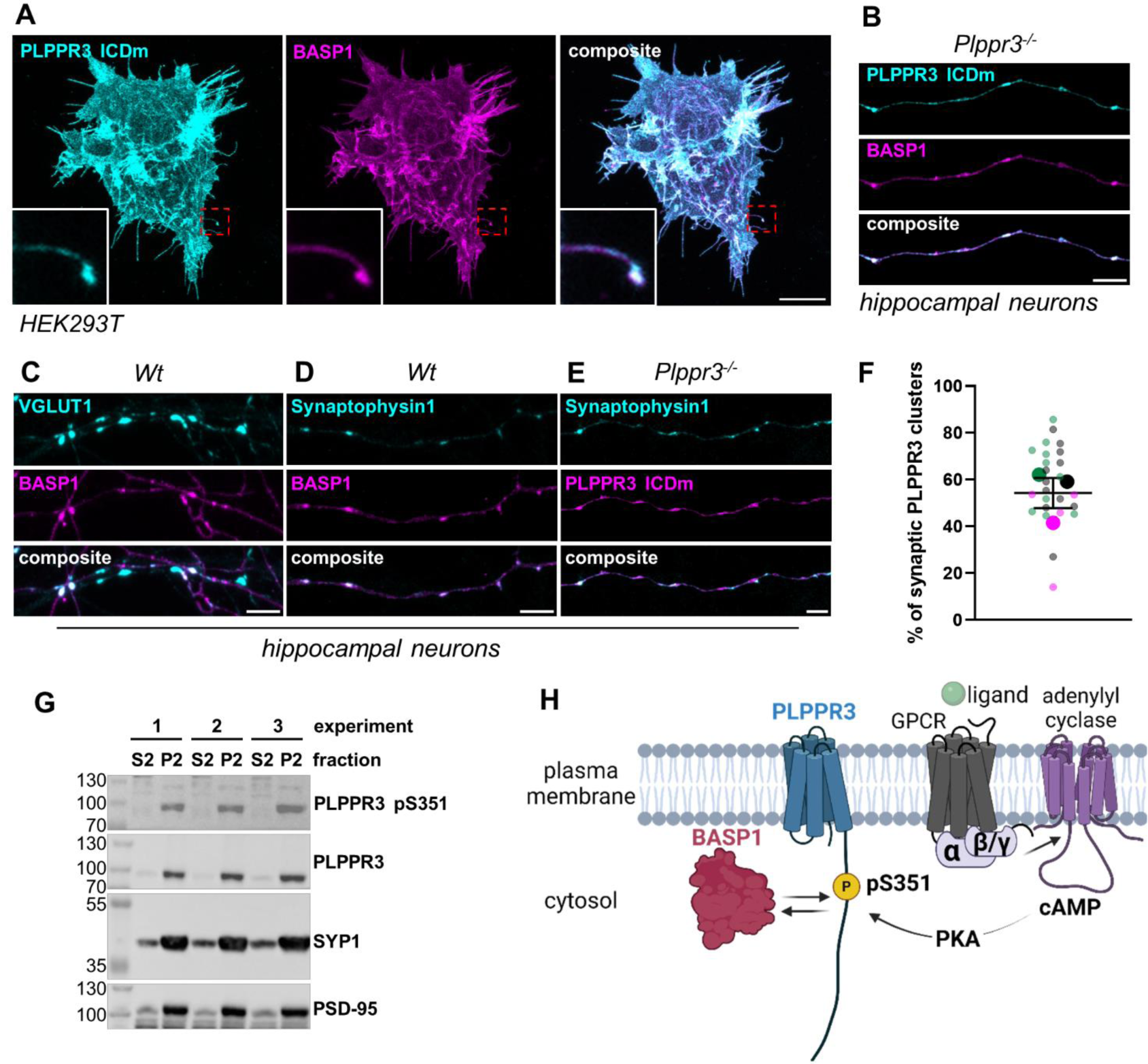
Characterization of presynaptic PLPPR3 ICDm-BASP1 complex. (A) Co-localization of ICDm and BASP1 in non-neuronal cells. HEK293T cells were fixed 12 hours after transfection, labelled and imaged. Red boxes indicate zoomed in areas. Scale bar = 10 µm. (B) Co-localization of ICDm and BASP1 in primary hippocampal neurons. (C) Co-localization of endogenous VGLUT1 and BASP1 in primary hippocampal neurons. (D) Co-localization of Synaptophysin1 and BASP1 in primary hippocampal neurons. (E) Co-localization of Synaptophysin-1 and PLPPR3 ICDm in primary hippocampal neurons. (F) Quantification of PLPPR3 ICDm clusters inside synapses (mean ± SEM). N=3, n=5-11. In (B)-(E), neurons were transfected with respective recombinant proteins at DIV1 and analyzed at DIV7. Scale bar = 5 µm. (G) Localization of PLPPR3 and pS351 to synaptosomal fractions. Crude synaptosomes were prepared from adult mouse brain and analyzed by western blot. S2 = cytosolic fraction, P2 = crude synaptosomes, SYP1 = synaptophysin1. (H) Working model of presynaptic PLPPR3-BASP1 complex. Created with Biorender.com.

## Discussion

In summary, the work presented here implicates PLPPR3 membrane proteins as signaling integrators at neuronal synapses. Our work demonstrates that the PLPPR3 intracellular domain is highly phosphorylated, and that one of the residues – S351 – is a *bona fide* phosphorylation site of PKA. Phosphorylation of PLPPR3 at S351 regulates binding to BASP1, a protein that has previously been implicated to play key roles during neuronal morphogenesis and regeneration.^35,36^ BASP1 and PLPPR3 intracellular domain co-localize in presynaptic compartments, and PLPPR3 and PLPPR3 pS351 are exclusively enriched in synaptosomes isolated from adult mouse forebrain. Thus, the work presented here describes a novel protein complex in the presynaptic compartment with hitherto unknown function, and a PKA signaling event that regulates their binding.

### The intracellular domain of PLPPR3 is highly phosphorylated

Our work identifies 26 high-confidence phosphorylation sites in the 433 amino acid-long PLPPR3 intracellular domain, which form two dense clusters. One of them stretches from amino acids 343-400 and includes on average one phosphorylation site every five amino acids. The second cluster stretches from amino acids 560-575, where on average every third residue is phosphorylated (see Figure 2C). Curiously, both of these phosphorylation clusters overlap with the few regions in the intracellular domain that are well conserved between the five PLPPR family members. This is interesting, because PLPPR intracellular domains are of significantly different lengths, ranging from ∼50 to ∼400 amino acids, and generally poorly conserved. The first phosphorylation cluster overlaps with the region of hydrophobic and positively charged amino acids (aa 341-352 in PLPPR3), while the second phosphorylation cluster overlaps with a region of phosphorylatable and negatively charged residues (aa 555-567).^1^ Thus, these regions could be important regulatory domains in all PLPPRs.

While these phosphorylation hotspots show significant overlap with conserved regions, few of the actual phosphorylation sites are well conserved. Some of the regulatory sites that stand out are PLPPR3 S343, which is present in PLPPR1 and PLPPR4, while in PLPPR5 this residue is substituted with the negatively charged glutamic acid, and S565, which shows a conservative substitution to threonine in PLPPR4, and to negatively charged glutamic acid in PLPPR1 and PLPPR5. Thus, it is tempting to speculate that these sites could regulate functions common to all PLPPRs, such as filopodia formation. In contrast, other sites within these phosphorylation clusters and conserved regions – such as PLPPR3 S351 – are much less conserved among different PLPPRs, while showing high conservation in PLPPR3 from different evolutionary species (Figure 2B). Thus, it is likely that these sites regulate functions unique to PLPPR3. This is in good agreement with our data showing that S351 does not regulate filopodia formation (Figure 3H, I).

What could be the function of such a large number of phosphorylation sites? It is particularly interesting that the polyE box consisting of a stretch of 20 negatively charged glutamic acids is located between the two densely phosphorylated regions. It is tempting to speculate that these regions, together with the polyE box, could regulate the interactions of specific domains with the inner plasma membrane. Indeed, our unpublished data indicates that PLPPR3 binds membrane phosphoinositides via its intracellular domain, and the same has been published for PLPPR5.^11^ Thus, phosphorylation of specific domains could repel the intracellular domain from the inner plasma membrane, abolishing interactions with anionic membrane lipids and, simultaneously, making it available for protein-protein interactions. Phosphorylation is indeed a common way for cells to regulate protein interactions, and computational analysis shows that binding regions within a protein have a tendency to be phosphorylated.^33^ This suggests a similarity to known presynaptic scaffold proteins bassoon and piccolo, which have 48 and 31 phosphorylation sites, respectively.^38^ Besides interactions, a high number of phosphorylation sites have been reported for ion channels, where they fine-tune channel gating and thereby neuronal activity.^39–41^ Finally, a large number of phosphorylation sites suggest a convergence of signaling pathways on the intracellular domain of PLPPR3. Indeed, our preliminary data indicate that GSK3 can phosphorylate PLPPR3, but prediction tools suggest many other kinases are also involved (data not shown). Taken together, our data positions PLPPR3 intracellular domain as an intriguing signaling switch at the interface of the cell membrane and cytosol.

### PLPPR3 is a synaptic protein phosphorylated by PKA at S351

One of the phosphorylation sites that we identified in our mass spectrometry approach was S351 (Figure 2B). This site has also previously been identified in other phosphoproteome studies using brain tissue.^42–44^ Our data indicates that there are low steady-state levels of S351 phosphorylation at all developmental stages in primary neurons (Figure 4A), as well as in adult brain tissue (Figure 4C). This is in agreement with mass spectrometry studies showing that PLPPR3 S351 is phosphorylated in neonatal as well as 3-week-old mouse brain.^43,44^

Our data indicates that PLPPR3 S351 is a functionally relevant site. This idea is supported by the fact that this site is evolutionarily fully conserved between PLPPR3 from different species (Figure 2B) and is partially also conserved in other family members (see discussion above). Furthermore, S351 phosphorylation can be triggered in primary neurons (Figure 4B) as well as in brain tissue (Figure 4C). A recent phosphoproteomic study indicates that PLPPR3 S351 phosphorylation is also regulated endogenously *in vivo*. Pinto et al. (2022) studied how microglia control sleep homeostasis by releasing TNFα, a signaling molecule that controls sleep. This regulates the phosphorylation of thousands of proteins in the cortex during sleep, and curiously, among others, dynamic regulation of PLPPR3 pS351 was observed.^45^ Thus, it appears that PLPPR3 S351 is a functionally relevant site, and it would be worth testing its function in the context of sleep.

We rigorously characterized the phosphorylation of PLPPR3 S351 by PKA in a cAMP-dependent manner (Figures 3A-D) and discovered that PKA targets S351 in the PLPPR3 intracellular domain (Figures 3E-F). While the best-studied function of PLPPR3 is filopodia formation, and PKA has been shown to regulate filopodia formation,^18^ phosphorylation of S351 does not appear to regulate this function (Figures 3H, I). Instead, we found that phosphorylation of PLPPR3 regulates binding to BASP1 (Figures 4D-F) and that the two proteins co-localize in presynaptic clusters (Figures 5B-F). BASP1 is primarily known as a growth-associated protein that is highly expressed during early development and regeneration where it regulates axon growth.^35,36,46^ The mechanism is thought to involve regulation of the actin cytoskeleton.^34,35^ However, several low- and high-throughput studies have suggested a synaptic function for BASP1. Early electron microscopy studies identified BASP1 localization to presynaptic densities and synaptic vesicles.^47,48^ Immunolabeling experiments demonstrated the co-localization of BASP1 with the presynaptic protein VAMP2 in DIV12 primary hippocampal neurons as well as adult cerebellar tissue.^49^ Additionally, discovery mass spectrometry studies consistently find BASP1 in the synaptic fractions.^50–54^ Nevertheless, to date, the function of BASP1 in synapses has not been characterized.

Our experiments are the first to demonstrate the localization of PLPPR3 to presynapses. In early development, PLPPR3 localises to axons, where it is most highly expressed during the time when axons undergo branching.^3^ It appears that beyond the peaks of the branching stage, PLPPR3 enriches in immature presynapses, as demonstrated by co-localization with Synaptophysin-1 (Figures 5E,F). Furthermore, we have recently established that PLPPR3 is localized in synaptosomes prepared from different adult brain areas (Figure 5G).^5^ Importantly, we also found PLPPR3 pS351 exclusively in synaptosomal fractions (Figure 5G). Adding further evidence, phosphoproteome studies consistently find PLPPR3 pS351 in synaptic fractions.^45,55–58^ Taken together, these data indicate that PLPPR3 is a synaptic protein. It is tempting to speculate that phosphorylation may be the regulatory switch between different PLPPR3 functions. To this end, it is noteworthy that phosphorylation of endogenous PLPPR3 S351 was most potently triggered at 9 days *in vitro* (Figure 4B), which is precisely when cultured neurons undergo synaptogenesis (generally considered 7-14 days *in vitro*).^59^ Thus, it would be interesting to test whether phosphorylation of S351 – perhaps via interaction with BASP1 – regulates its accumulation at nascent presynaptic sites.

### Molecular mechanism and function of PLPPR3-BASP1 complex at the presynapse

PKA is a well-known modulator of both pre- and postsynaptic proteins, and it is considered one of the important players in learning and memory.^60,61^ A wealth of data connects PKA to synaptic transmission via presynaptic effectors such as Synapsin-1, Synaptotagmin-12 and Rim1 to name a few.^62–64^ Recent evidence links PKA activity to synapse formation.^65,66^ What could be the function of PKA-induced PLPPR3-BASP1 complex at the synapse?

As both PLPPR3 and BASP1 interact with the plasma membrane and engage the actin cytoskeleton, the most likely mechanism of this protein complex at the presynapse is mediating the signaling between the membrane and the active zone. Accumulation of actin is one of the early steps in synaptogenesis, as it provides a scaffold for synaptic vesicles.^67,68^ Thus, presynaptic actin regulates neurotransmitter release and plays an important role in synaptic vesicle recycling.^69^ PKA has been described to regulate some of these functions.^70,71^ In this light, as the expression profiles of both PLPPR3 and BASP1 are higher during early development, it would be interesting to test whether this protein complex may play a role in synapse formation by engaging the actin cytoskeleton. On the other hand, BASP1 has several interaction partners – the kainate receptor GluK1, and the synaptic vesicle associated proteins synaptojanin-1 and dynamin-1 – that, together with its reported localization to synaptic vesicles (see discussion above) could also suggest a role in synaptic transmission or synaptic vesicle recycling.^72–74^ Future studies should thus focus on understanding the role of PLPPR3-BASP1 complex at presynapses.

Finally, what could be the signaling pathway that controls the presynaptic interaction of PLPPR3 and BASP1? The activity of adenylyl cyclases, which produce cAMP essential for the activation of PKA, is controlled by GPCRs, of which the Gi/o -coupled receptors inhibit, and Gs-coupled receptors activate cAMP production. To date, many well-studied neuromodulators have been shown to regulate PKA signaling. Adenosine, serotonin, endocannabinoids, dopamine as well as vasoactive intestinal protein (VIP) all regulate PKA via various receptors in the brain.^75–81^ Future studies could investigate which extracellular signal regulates the phosphorylation levels of PLPPR3 S351 with GPCR subtype-specific ligands.

### Conclusion

The work presented here establishes PLPPR3 as a synaptically-enriched protein that is highly phosphorylated. Through its localization to the plasma membrane and long intracellular domain, it is ideally suited to integrate extra-and intracellular signals and mediate responses. Via PKA-regulated phosphorylation of S351, PLPPR3 binds BASP1, and the protein complex is enriched at the presynapse, where it may play a role in synaptic maintenance or transmission. Future studies will focus on elucidating the role of this novel protein complex at the synapse.

## Materials and Methods

### Antibodies

#### Animal procedures

Animals were handled and housed in accordance with the local ethical guidelines and regulations. The animals were kept under standard conditions with 12-hour light/dark cycle and food and water available at all times. All animal experiments were registered in the Landesamt für Gesundheit und Soziales (LaGeSo) under license T0347/11. All animals used in this work were with the C57 Bl/6NCrl genetic background. Primary cultures from wild-type and *PTEN^fl/fl^* animals were used as control for the *Plppr3^-/-^*.^3,82^ No experiments were analyzed by sex.

#### Cell line culturing and transfection

HEK293T (#LV900A-1, BioCAT/SBI, RRID: CVCL_UL49) and N1E-115 (#CRL-2263, ATCC, RRID: CVCL_0451) cells were cultured in DMEM high glucose (#11965092, Life Technologies) supplemented with 10% fetal bovine serum (#F7524, Sigma; heat inactivated) and 1% penicillin/streptomycin (#15140122, Gibco) at 37°C with 5% CO_2_. Cells were passaged twice a week at a ratio of 1:4.

HEK293T cells were plated on poly-DL-ornithine (#P8638, Sigma; 15 μg/ml for 1 hour at 37°C) coated glass coverslips or plastic cell culture dishes. N1E-115 cells were grown without substrate. For western blot and immunoprecipitation experiments, cells were plated at a density of 300 000/well on 6-well culture dishes (#92006, TPP). For immunocytochemistry experiments, cells were plated at a density of 20 000 (HEK293T) or 30 000 (N1E-115) per well on 12-well culture plates (#92012, TPP). Cells were grown overnight before transfection.

Cell lines were transfected with Lipofectamine 2000 (#11668019, Invitrogen) in Opti-MEM reduced serum medium (#31985062, Gibco), always using 1 μg of total DNA and 2 μl of Lipofectamine per well. For 12-wells, 300 μl of Opti-MEM/well was used, for 6-well plates 400 μl/well was used. Transfection mix was prepared in two steps. First, half of the Opti-MEM volume was mixed with DNA, and the other half with Lipofectamine, mixed, and incubated at room temperature for 5 minutes. Then, the DNA and Lipofectamine were pooled and incubated on the water bath at 37°C for 10 minutes. Transfection mix was added dropwise directly to the cell culture medium and incubated for 5 hours. Finally, the medium was aspirated and exchanged with fresh complete medium. For experiments that required starvation, cells were changed into FBS-free medium consisting only of DMEM and penicillin/streptomycin. Cells were grown overnight before experimental procedures.

#### Primary neuronal culture and transfection

To prepare primary neuronal cultures, cortices and hippocampi of embryos (both female and male) from E16.5 pregnant mice were dissected into individual Eppendorf tubes containing cold HBSS (2 hippocampi/tube, one hemisphere of cortex/tube). The tissue was rinsed with fresh HBSS and digested with 20 U/ml papain (#LS003126, Worthington Biochemical) (500 μl per one hippocampi Eppendorf, 1 ml for cortex) for 30 minutes on thermomixer at 37°C with gentle shaking (350 rpm). Papain digestion was stopped by replacing the supernatant with prewarmed inactivation solution [DMEM with 10% FCS, 1% Pen/Strep, 2.5 mg/ml Albumin/BSA (#A2153, Sigma) and 2.5 mg/ml trypsin inhibitor (#T9253, Sigma)] and incubating at 37°C for 5 minutes. Then, supernatant was replaced with 200 μl plating medium (Neurobasal, 10% heat inactivated FCS, 2% B27, 1% GlutaMax and 1% Pen/Strep) and tissue was disrupted by gentle trituration. Neurons from different preparation tubes were pooled and counted.

Neurons were plated in plating medium and after 2 hours changed to complete medium (Neurobasal, 2% B27, 0.5% Glutamax and 1% Pen/strep). For western blot experiments, cortical or hippocampal neurons were plated on poly-DL-ornithine (15 μg/ml) coated plastic 6-well dishes at a density of 500 000/well. For immunocytochemistry experiments, hippocampal neurons were plated on poly-DL-ornithine (15 μg/ml) + laminin (20 μg/ml; #L2020, Sigma-Aldrich) coated 18 mm glass coverslips at a density of 120 000/well. Primary cultures were grown under constant culture conditions (37°C, 5% CO_2_) until further use.

For immunocytochemistry experiments on figure 5B-F, hippocampal neurons were transfected at DIV1 using the calcium phosphate method.^83^ In total, 4 μg of DNA was used per well (2+2 μg/well for double transfections). To prepare the transfection solution, CaCl_2_ (2.5 μl/well) was diluted in H_2_O (22.5 μl/well) and briefly mixed, then DNA was added and mixed. Then, 2xBSS buffer (25 μl/well; 50 mM BES, 280 mM NaCl, 1.5 mM Na_2_HPO_4_, pH 7.26) was added dropwise and the solution was gently mixed by tapping. Finally, pre-warmed Neurobasal medium with 2% B27 and 0.25% GlutaMax (450 μl/well) was added, and the transfection mix was incubated for 15 minutes at 37°C. Conditioned culture medium was collected from each well and exchanged with transfection medium, and neurons were incubated for 20 minutes. Each well was washed three times with 500 μl warm HBSS buffer (135 mM NaCl, 4 mM KCl, 1 mM Na_2_HPO_4_, 2 mM CaCl_2_, 1 mM MgCl_2_, 20 mM HEPES, 20 mM d-Glucose, pH 7.3). After the final wash, warm conditioned medium was added back to wells. Neurons were grown until further use.

#### Acute brain slice preparation and stimulation with Forskolin

*Wt* C57/BL6 mice were deeply anesthetized with isoflurane and then decapitated. The brains were quickly cooled in ice cold solution (110 mM choline chloride, 2.5 mM KCl, 1.25 mM NaH2PO_4_, 26 mM NaHCO_3_, 11.6 mM sodium ascorbate, 3.1 mM sodium pyruvate, 7 mM MgCl_2_, 0.5 mM CaCl_2_ and 10 mM d-glucose, pH 7.4) and sectioned into 300 μm thick coronal slices on a vibrotome. The acute brain slices were then incubated in the same solution at 32°C for 5 minutes, followed by incubation in artificial cerebrospinal fluid (ACSF; 125 mM NaCl, 2.5 mM KCl, 1.25 mM NaH_2_PO_4_, 25 mM NaHCO_3_, 2 mM CaCl_2_, 1 mM MgCl_2_, and 25 mM dextrose) at 32°C for 30 minutes prior to use. Each brain slice was further dissected, and three half slices were used for each experimental condition. Test slices were placed into an incubation chamber containing ACSF with Forskolin (30 μM) for 15 minutes at 32°C. Control slices remained in an incubation chamber with ACSF lacking Forskolin. Following Forskolin stimulation, test and control brain slices were placed in separate Eppendorf tubes and snap frozen in liquid nitrogen and stored at -80°C until further use. All solutions were saturated with 95% O_2_/5% CO_2_.

### Cloning

All plasmids generated for this work are in pCAX backbone (CAG promoter) and have a C-terminal tag (Flag tag separated with a flexible GSGGGSG-linker, His tag with a VDGRP-linker). All constructs were validated by control digest and sequencing. A detailed protocol that describes the generation of all the plasmids used in this work can be found at dx.doi.org/10.17504/protocols.io.dm6gp37j8vzp/v1.

### Drug treatments: DMSO, Forskolin, H89, 8-Br-cAMP

All drugs were added directly into cell culture medium and mixed by gentle trituration. For acute brain slice treatment, Forskolin was added into 100 ml ACSF, and the mixture was allowed to homogenize by air flow for a few minutes before brain slices were added. For experiments requiring DMSO control treatments, at least one control treatment was carried out per experiment to ensure the solvent has no effect. Forskolin (#11018, Cayman Chemical) was used at a final concentration of 30 μM for 5-15 minutes, depending on the experiment (see figure descriptions). 8-Br-cAMP (#B7880, Sigma-Aldrich) was used at 0.5-1 mM concentration for 5 minutes. H89 (#2910, Tocris) was used at 1-50 μM concentration for 1 hour prior to Forskolin treatment.

### Cell and tissue lysis

To lyse cells and neurons, culture medium was aspirated, and cells were rinsed with ice cold PBS (prepared from tablets, #A9191, Applichem). PBS was aspirated and lysis buffer (200μl for neurons, 300 μl for cell lines) was added, cells were scraped off the surface and collected into an Eppendorf tube. Cells were incubated in lysis buffer at 4°C for 20 minutes with overhead rotation, followed by centrifugation at 14 000 rpm for 20 minutes at 4°C. Supernatant was collected into a fresh tube, and samples were either used directly or kept in -20°C until use.

To prepare brain lysates used in figure 4D, whole adult mouse brain (ca 0.45 g) was mechanically disrupted in 4.5 ml lysis buffer using Dounce homogenizer. To prepare brain slice lysates used in figure 4C, the slices were homogenized in 1 ml lysis buffer using micro tissue homogenizer and trituration with pipet. In both cases, tissue homogenates were incubated in lysis buffer at 4°C for 20 minutes with overhead rotation and centrifuged at 14 000 rpm for 20 minutes at 4°C. Supernatant was collected into a fresh tube, and samples were either used directly or kept in -20°C until use.

All samples were lysed in house-made Ripa buffer (50 mM Tris-HCl, 150 mM sodium chloride, 1% NP-40, 0.5% sodium deoxycholate and 0.1% sodium dodecyl sulfate, pH 8.0) supplemented with lab-made phosphatase inhibitors [1 mM Na2MO4, 1 mM NaF, 20 mM β-glycerophosphate, 1 mM Na3VO4 and 500 nM Cantharidin (#3322.1 Roth)] and commercial protease inhibitors (protease inhibitor cocktail III, #539134, Calbiochem). Phosphatase inhibitors were omitted when Lambda phosphatase treatment was required. All samples used for western blot experiments, except the co-immunoprecipitation samples, were measured for total protein concentration using the Pierce BCA Protein Assay Kit (#23225, Thermo Scientific) following the manufactureŕs protocol.

### Lambda phosphatase treatment

48 μl of neuron or cell lysate was mixed with 6 μl of 10x NEBuffer for Protein MetalloPhosphatases (PMP), 6 μl of 10 mM MnCl and 6 μl (=2400 units) of Lambda phosphatase (#P0753, NEB). Dephosphorylation was carried out at 30°C for 30 minutes and stopped by addition of Roti Load buffer (#K929.1, Roth). Treatment control included H_2_O instead of Lambda phosphatase and was also incubated at 30°C for 30 minutes. Phosphorylated control was diluted with H_2_O to the same final volume of other samples and kept on ice for 30 minutes.

### *In vitro* phosphorylation assay

*In vitro* phosphorylation assay was carried out using purified PLPPR3 ICD from E.coli. Prior to phosphorylation assay, protease inhibitors were added to the purified protein (1:100 AEBSF; #1421, Applichem). 4.1 μl of PLPPR3 ICD (final concentration 0.075 mg/ml) was mixed with 0.2 μl purified PKA catalytic subunit (final concentration 20 000 Units; #P6000S, Biolabs), 0.8 μl phosphorylation buffer (final concentration of 25 mM HEPES, 100 mM NaCl, 5 mM MgCl, 2 mM EGTA, 25 mM DTT), 1 μl ATP (final concentration 1 mM) and 14 μl H_2_O. Phosphorylation was carried out at 30°C for 2 hours. The reaction was stopped by the addition of Roti Load buffer.

### Co-immunoprecipitation

Cell lysates (prepared as described above) from 2 wells of 300 000 HEK293T cells each were pooled. 30 μl of protein lysate was taken for input control, mixed with Roti Load and boiled for 5 minutes at 95°C .12 μl of tGFP or IgG antibody (roughly 1:50) was added to the rest of the protein lysate and incubated at 4°C overnight with overhead rotation. Dynabeads Protein A (#10002D, Invitrogen) were washed with 1 ml Ripa buffer three times, and after the final wash reconstituted in their original volume with Ripa. 20 μl of bead slurry was added to each sample. The sample was incubated for 1 hour at 4°C with overhead rotation and washed 4 x 15 minutes with Ripa at 4°C with overhead rotation. Finally, the beads were eluted with 30 μl 1x Roti Load by boiling at 95°C for 3 minutes. This step was repeated twice to maximize yield.

### Preparation of crude synaptosomes

Crude synaptosomes were prepared from the brains of 5-week-old female and male mice. Brains were dissected to exclude the cerebellum and the olfactory bulb. The brain was weighed and homogenized in 1 ml/100 mg homogenization buffer (final concentration 5 mM Tris, 1 mM EDTA, 0.32 M sucrose, pH 7.4) supplemented with protease and phosphatase inhibitors (see section on Cell and tissue lysis). The lysate was cleared of nuclei and cell debris by centrifugation at 1000 g for 10 minutes. Supernatant (S1) was collected and centrifuged at 15 000 g for 30 minutes, after which the supernatant (S2) was collected. The remaining pellet containing the synaptosomes (with myelin, membranes and mitochondria) was resuspended in homogenization buffer without sucrose (5 mM Tris, 1 mM EDTA, pH 7.4) and centrifuged at 15 000 g for 30 minutes. Supernatant was discarded and the pellet (P2) was snap frozen. All work was carried out at 4°C or on ice.

Prior to western blot analysis, S2 and P2 fractions were thawn on ice, and the P2 were resuspended in 150 μl homogenization buffer (roughly equal to the volume of S2 fractions). The samples were incubated on ice with 15 μl of 10% TritonX-100 for 10 minutes to break open synaptosomal membranes. Total protein concentration was measured with BCA assay (see Cell and tissue lysis) and 25 μg of total protein was loaded on the gel.

### SDS-PAGE, PhosTag SDS-PAGE and western blotting

We used 20 μg of total protein from N1E-115 cells or 40 μg of total protein from neuronal and tissue lysates for SDS-PAGE western blot analysis. Protein lysates were separated on 8-12% acrylamide gels at 80V for 15 minutes followed by 120V until the dye front ran out. Proteins were transferred onto 20 μm pore size nitrocellulose membranes (#1620097, Biorad) for 2.5 hours at 400 mA on ice. Membrane was briefly rinsed in dH_2_O, labeled with Ponceau S (#A2935, Applichem) solution according to manufactureŕs protocol and de-stained in dH_2_O. The membrane was blocked in 5% skim milk (#T145.2, Carl Roth) in TBS-T (5 mM Tris-HCl, 15 mM NaCl, 0.005% Tween20, pH 7.4) for 1 hour at room temperature and incubated with primary antibodies (see Table 1) in blocking solution at 4°C overnight. Membrane was washed 3 x 10 minutes in TBS-T and incubated with secondary antibodies (see Table 1) in blocking solution at room temperature for 1 hour, followed by 3 x 10 minutes washes in TBS-T. Blots were developed with ECL Western Blotting Substrate (#W1001, Promega) or ECL Select Western Blotting Detection Reagent (#RPN2235, Cytiva) according to manufactureŕs protocol. Images of western blots were acquired with Fusion SL camera (VilberLourmat, Germany) and manufactureŕs software using automatic exposure mode. Chemiluminescent signal image and molecular weight marker image were automatically overlaid by the software. For all SDS-PAGE experiments presented in this work, molecular weight marker ranging from 10 to 250 kDa was used (#26620, Thermo Scientific).

**Table 1.**
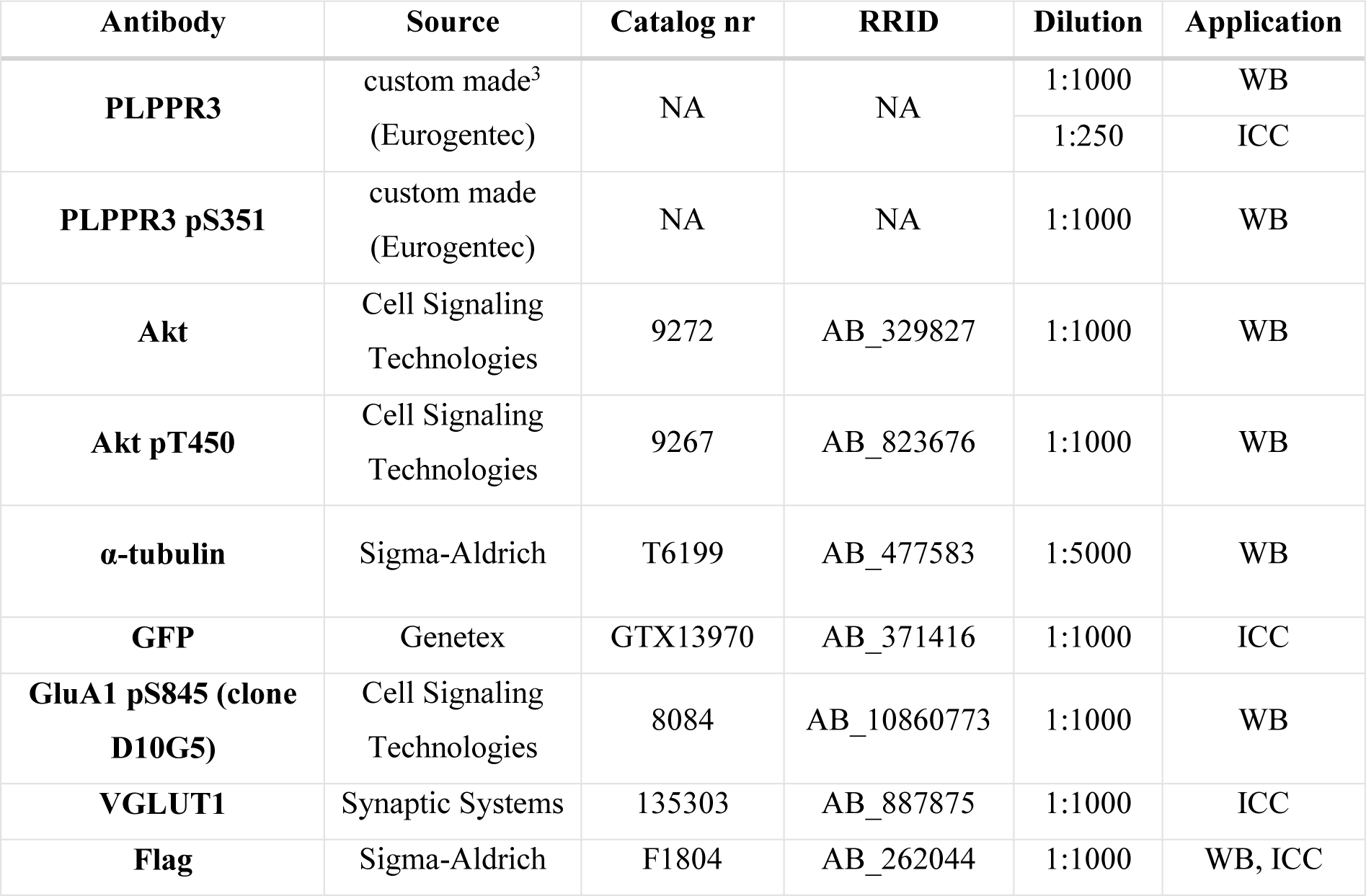
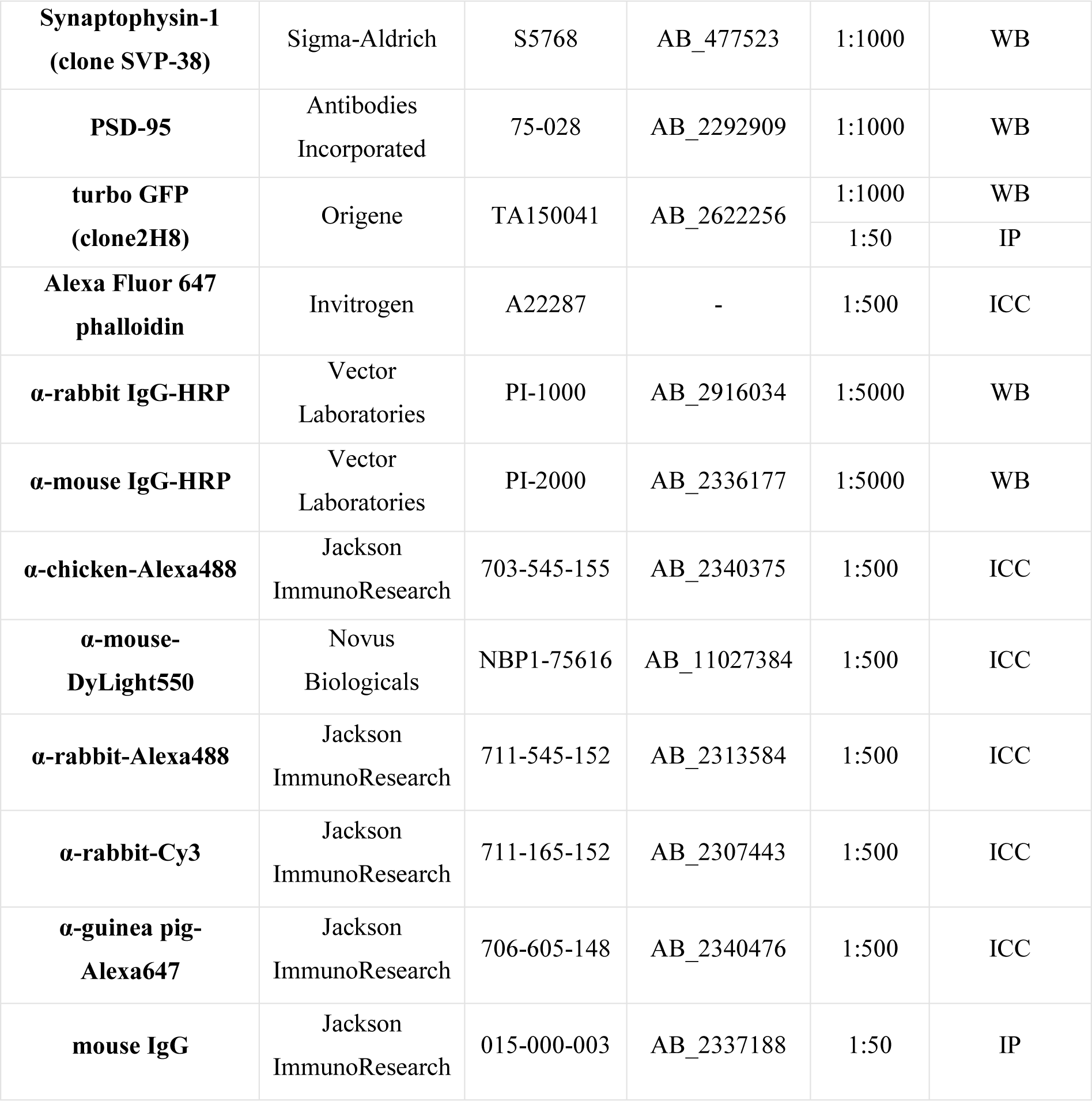
Antibodies used in this work. NA – not applicable, WB – western blot, ICC – immunocytochemistry, IP – immunoprecipitation.

| Antibody | Source | Catalog nr | RRID | Dilution | Application |
| --- | --- | --- | --- | --- | --- |
| **PLPPR3** | custom made<sup>3</sup><br>(Eurogentec) | NA | NA | 1:1000 | WB |
|  |  |  |  | 1:250 | ICC |
| **PLPPR3 pS351** | custom made<br>(Eurogentec) | NA | NA | 1:1000 | WB |
| **Akt** | Cell Signaling<br>Technologies | 9272 | AB_329827 | 1:1000 | WB |
| **Akt pT450** | Cell Signaling<br>Technologies | 9267 | AB_823676 | 1:1000 | WB |
| **α-tubulin** | Sigma-Aldrich | T6199 | AB_477583 | 1:5000 | WB |
| **GFP** | Genetex | GTX13970 | AB_371416 | 1:1000 | ICC |
| **GluA1 pS845 (clone D10G5)** | Cell Signaling<br>Technologies | 8084 | AB_10860773 | 1:1000 | WB |
| **VGLUT1** | Synaptic Systems | 135303 | AB_887875 | 1:1000 | ICC |
| **Flag** | Sigma-Aldrich | F1804 | AB_262044 | 1:1000 | WB, ICC |
| **Synaptophysin-1**<br>**(clone SVP-38)** | Sigma-Aldrich | S5768 | AB_477523 | 1:1000 | WB |
| **PSD-95** | Antibodies<br>Incorporated | 75-028 | AB_2292909 | 1:1000 | WB |
| **turbo GFP**<br>**(clone2H8)** | Origene | TA150041 | AB_2622256 | 1:1000 | WB |
|  |  |  |  | 1:50 | IP |
| **Alexa Fluor 647**<br>**phalloidin** | Invitrogen | A22287 | - | 1:500 | ICC |
| **α-rabbit IgG-HRP** | Vector<br>Laboratories | PI-1000 | AB_2916034 | 1:5000 | WB |
| **α-mouse IgG-HRP** | Vector<br>Laboratories | PI-2000 | AB_2336177 | 1:5000 | WB |
| **α-chicken-Alexa488** | Jackson<br>ImmunoResearch | 703-545-155 | AB_2340375 | 1:500 | ICC |
| **α-mouse-**<br>**DyLight550** | Novus<br>Biologicals | NBP1-75616 | AB_11027384 | 1:500 | ICC |
| **α-rabbit-Alexa488** | Jackson<br>ImmunoResearch | 711-545-152 | AB_2313584 | 1:500 | ICC |
| **α-rabbit-Cy3** | Jackson<br>ImmunoResearch | 711-165-152 | AB_2307443 | 1:500 | ICC |
| **α-guinea pig-**<br>**Alexa647** | Jackson<br>ImmunoResearch | 706-605-148 | AB_2340476 | 1:500 | ICC |
| **mouse IgG** | Jackson<br>ImmunoResearch | 015-000-003 | AB_2337188 | 1:50 | IP |

**Table 2.**
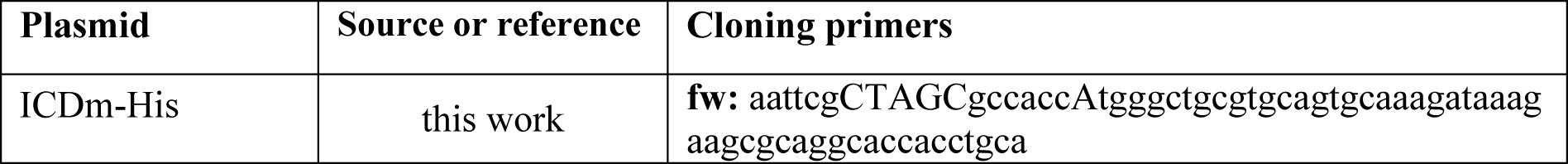
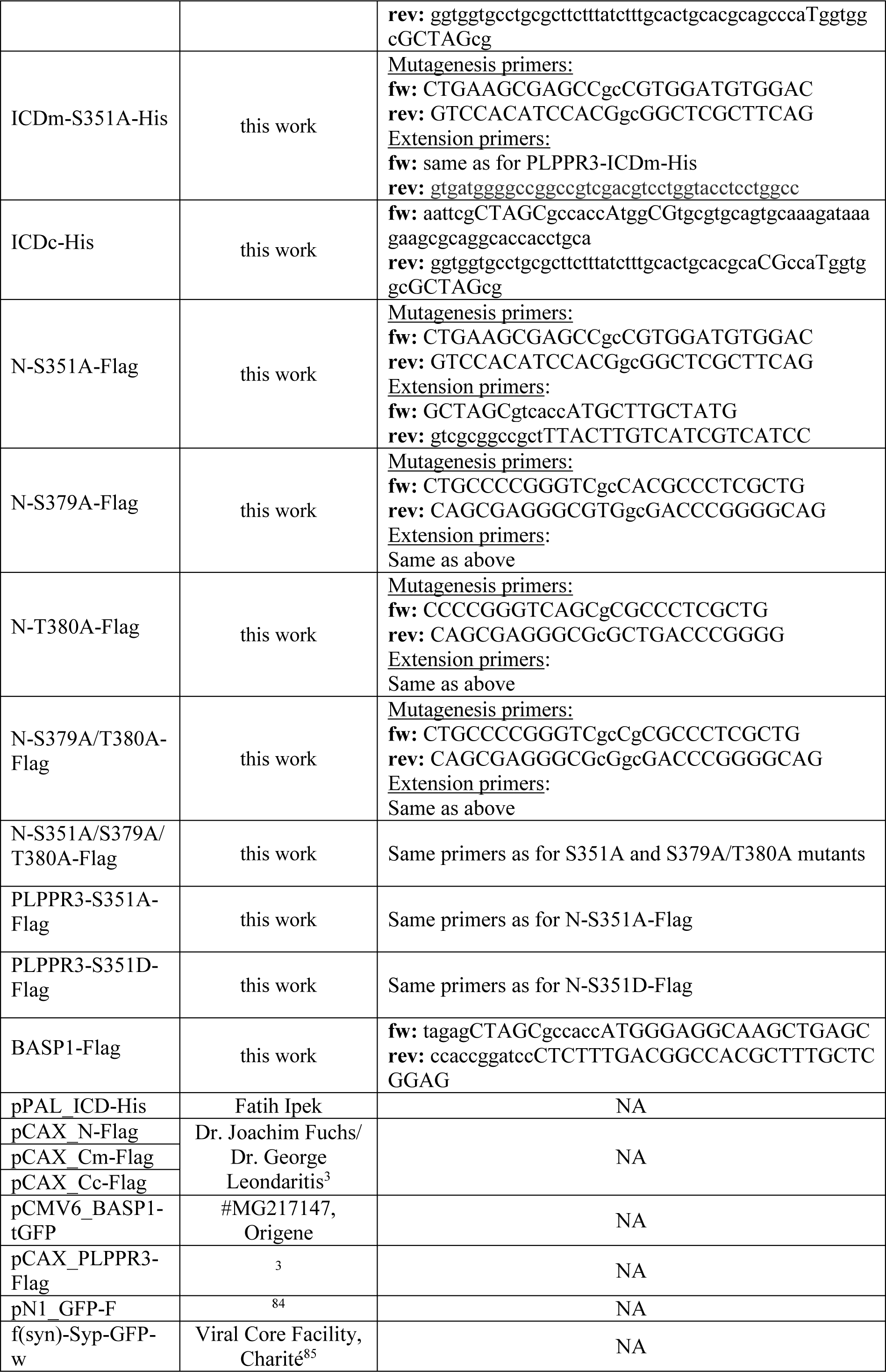
Plasmids used in this work. ICD – intracellular domain, Fw – forward, rev – reverse, NA – not applicable.

For phosphorylation analysis, 8% 50 μM Zn^2+^ PhosTag acrylamide (#AAL-107, FUJIFILM) gels were prepared according to manufactureŕs protocol. 40 μg of total protein was loaded and gel electrophoresis was run under a constant current (30 mA/gel) until the dye front ran out. Gels were washed 3 x 10 minutes in transfer buffer containing 10 mmol/L EDTA, and 1 x 10 minutes in transfer buffer without EDTA prior to transfer. Proteins were transferred onto a PVDF membrane with 0.2 μm pore size (#88520, Thermo Scientific) for 2.5 hours at 400 mA on ice. The rest of the steps were carried out as described above. PhosTag blots do not contain molecular weight markers as they do not give meaningful information about the size of proteins and may cause distortion of protein bands (see PhosTag SDS-PAGE guidebook, available at http://www.bujnochem.com/wp-content/uploads/2019/09/FUJIFILM-Wako_Phos-tag-R.pdf). All western blot experiments, with the exception of Figure 4A-C, were performed a minimum of three times with different passages of cells.

### Quantification of western blot band intensities

Quantification of protein band intensities was carried out in Fiji using the Analyze->Gels command. Composed protein band and molecular weight marker images were used for quantification. Rectangular selection, big enough to include the biggest band, was used to select each protein band. Each lane was plotted using the Plot lanes command, and if necessary, a straight line was drawn to close off the area under the curve. Intensity was measured for each band. Co-immunoprecipitation band (PLPPR3) was normalized to immunoprecipitation band (tGFP) by dividing the intensity PLPPR3/tGFP. The value of each non-stimulated replicate was normalized to 1, and the value of Forskolin-stimulated sample was expressed as Forskolin/control.

### Purification of PLPPR3 ICD constructs

12 x 10^6^ HEK293T cells were seeded in 150 cm^2^ Corning flasks (#CLS431465, Merck) and cultured overnight. Cells were transfected with ICDm-His or ICDc-His constructs using 75 μg of total DNA and 225 μl PEI in 5 ml serum free hybridoma medium (#11279023, Gibco), and grown overnight. Cells were washed with ice cold PBS once and scraped off the flask in fresh 10 ml ice cold PBS on ice. Following centrifugation at 4000 rpm for 1 min at 4°C, PBS was aspirated, and the cell pellet was frozen.

Four independent replicates were purified and analyzed in parallel. Frozen pellets were thawn on ice, lysis buffer was added [20 mM Hepes, 150 mM NaCl, 20 mM Imidazole, phosphatase inhibitor cocktail (1:25), Cantharidin (1:50), protease inhibitor tablets (1 tbl/10 ml, #4693159001, Roche), AEBSF (1:100, #A1421, Applichem), pH 7.4] and the pellets were sonicated for 3 x 60 seconds with 6 cycles and 60% power. Lysate was cleared by centrifugation at 21 000 g at 4°C for 30 minutes. For the purification of cytosolic ICDc-His, the supernatant was collected and incubated with Ni SepharoseTM 6 Fast Flow (GE) beads (300 μl/sample) overnight at 4°C on a rotator. For the purification of membrane-bound ICDm-His, protein was extracted from the pellet with 0.5% Fos-choline 14 (#F312, Anatrace) at 4°C for 40 minutes and centrifuged at 21 000 g for 1 hour at 4°C. Supernatant was collected and incubated with Ni SepharoseTM 6 Fast Flow (GE) beads overnight at 4°C on a rotator. Sample was cleared on Pierce centrifugal columns (#89896, Thermo Scientific) by gravity flow and washed three times with 20 ml wash buffer (wash 1: 20 mM Hepes, 150 mM NaCl, 50 mM Imidazole, pH 8.5; wash 2: 20 mM Hepes, 500 mM NaCl, 50 mM Imidazole, pH 8.5; wash 3: 20 mM Hepes, 150 mM NaCl, pH 8.5). Finally, beads were solubilized in 800 μl of buffer (20 mM Hepes, 150 mM NaCl, pH 8.5). The buffer was discarded and dry beads with protein were handed to the mass spectrometry facility. Phosphorylation status of samples was confirmed by PhosTag SDS-PAGE prior to mass spectrometry analysis.

### Affinity chromatography for the identification of PLPPR3 pS351 interaction partners

Sulfolink columns (#44999, Thermo Scientific) were coupled with PLPPR3 peptides constituting the S351 phosphorylation site (phospho: LKRApSVDVDLLA; non-phospho: LKRASVDVDLLA) according to manufactureŕs protocol. Affinity purification was carried out following the manufactureŕs protocol using gravity flow instead of centrifugation. Affinity columns were equilibrated to room temperature for 15 minutes before use and equilibrated three times with 2 ml of PBS. Columns were incubated with 400 μl of freshly prepared brain lysate (see section on Cell and tissue lysis) in 1600 μl PBS for 1.5 hours at room temperature with overhead rotation. Columns were washed with 2 ml PBS four times and eluted with 2 ml of elution buffer (0.2 M glycine-HCl, pH 2.5) in ∼500 μl fractions into a fresh tube containing 100 μl neutralization buffer (1 M Tris-HCl, pH 8.5). The protein concentration in each fraction was measured by Nanodrop, and fractions with highest protein content were pooled to concentrate protein by ethanol precipitation. For EtOH precipitation, 9 volumes of ice cold 100% ethanol were added to 1 volume of protein solution. To aid precipitation, 2 μl of Glycoblue (#AM9515, Invitrogen) was added to the mix, and the protein was precipitated overnight at 4°C. The following day, the precipitation solution was centrifuged at 14 000 rpm for 1 hour at 4°C, the aqueous solution was discarded, and the protein pellet resolubilized in 100 μl Roti Load buffer and boiled at 95°C for 5 minutes. The protein samples were run on SDS-PAGE, pieces of gel were excised and analyzed by mass spectrometry. Four biological replicates consisting of brain lysates from four individual mice were performed.

### Phospho-mass spectrometry analysis

Prior to mass spectrometry analysis, beads were resuspended in 20 μl urea buffer (2M urea, 50 mM ammonium bicarbonate, pH 8.0), reduced in 12 mM dithiothreitol at 25°C for 30 minutes, followed by alkylation with 40 mM chloroacetamide at 25 °C for 20 minutes. Samples were digested with 0.5 μg Trypsin/LysC, Trypsin/GluC or GluC overnight at 30°C. The peptide-containing supernatant was collected, and digestion was stopped with 1% formic acid. Peptides were desalted and cleaned up using Stage Tip protocol (PMID: 12585499). Samples were eluted with 80% acetonitrile/0.1% formic acid, dried using speedvac, resuspended in 3% acetonitrile/0.1% formic acid and analysed by LC-MS/MS. Peptides were separated on a reversed-phase column, a 20 cm fritless silica microcolumn with an inner diameter of 75 μm, packed with ReproSil-Pur C18-AQ 3 μm resin (Dr. Maisch GmbH) using a 90 min gradient with a 250 nl/min flow rate of increasing Buffer B concentration (from 2% to 60%) on a High-Performance Liquid Chromatography (HPLC) system (Thermo Fisher Scientific), ionized with electrospray ionization (ESI) source (Thermo Fisher Scientific) and analyzed on a Thermo Q Exactive Plus instrument. The instrument was run in data dependent mode selecting the top 10 most intense ions in the MS full scans (ranging from 350 to 2000 m/z), using 70 K resolution with a 3×106 ion count target and 50 ms injection time. Tandem MS was performed at a resolution of 17.5 K. The MS2 ion count target was set to 5×104 with a maximum injection time of 250 ms. Only precursors with charge state 2–6 were selected for MS2. The dynamic exclusion duration was set to 30 s with a 10 ppm tolerance around the selected precursor and its isotopes. Raw data were analyzed using MaxQuant software package (v1.6.3.4, PMID: 27809316) using a human UniProt database (HUMAN.2019-07) and PLPPR3 sequence database, containing forward and reverse sequences. The search included variable modifications of serine, threonine, and tyrosine in addition to methionine oxidation, N-terminal acetylation, asparagine and glutamine deamidation. Carbamidomethylation of cysteine was set as a fixed modification. Minimal peptide length was set to seven amino acids and a maximum of 3 missed cleavages was allowed. The FDR was set to 1% for peptide and protein identifications. Phosphosite intensity values were normalized for PLPPR3 protein abundance. Results were filtered for reverse database hits and potential contaminants. Phosphosites found in 3/4 replicates with localization probability >0.75 and good MS2 spectra were considered high confidence.

### Mass spectrometry analysis for PLPPR3 pS351 interaction partners

The gel was digested with Trypsin prior to mass spectrometry analysis. Protein samples were concentrated on a trap column (PepMap C18, 5 mm x 300 μm x 5 μm, 100Ǻ, Thermo Fisher Scientific) with 2:98 (v/v) acetonitrile/water containing 0.1% (v/v) trifluoroacetic acid at a flow rate of 20 μl/ min for a total of 4 minutes. The samples were analyzed by nanoscale LC-MS/MS using a Q Exactive Plus mass spectrometer coupled with an Ultimate 3000 RSLCnano (Thermo Fisher Scientific). The system contained a 75 μm i.d. × 250 mm nano LC column (Acclaim PepMap C18, 2 μm; 100 Å; Thermo Fisher Scientific). The mobile phase A consisted of 0.1% (v/v) formic acid in H2O, while mobile phase B consisted of 80:20 (v/v) acetonitrile/H2O containing 0.1% (v/v) formic acid. The samples were eluted using a gradient of mobile phase B (3-53%) in 16 minutes, followed by washing with 98% of phase B and equilibration with starting condition using a flow rate of 300 nl/min. Full MS spectra (350–1,650 m/z) were acquired at a resolution of 70 000 (FWHM), followed by data-dependent MS/MS fragmentation (300-2,000 m/z) of the top 10 precursor ions (dissociation method HCD, resolution 17 500, 1+ charge state excluded, isolation window of 1.6 m/z, NCE of 27%, dd 10s, MS 1e6, MSMS AGC 5e5). Maximum ion injection time was set to 50 ms for MS, and 120 ms for MS/MS scans. Background ions at m/z 391.2843 and 445.1200 act as lock masses. Quantifiation of proteins was performed with MaxQuant software version 1.6.0.1 using default Andromeda LFQ parameter. Spectra were matched to murine (https://www.uniprot.org/; 17 073 reviewed entries), contaminant, and decoy database. The search included the following modifications: methionine oxidation and N-terminal acetylation (variable), carbamidomethyl cysteine (fixed). The false discovery rate was set to 0.01 for peptide and protein identifications. MS2 identifications were transferred between runs using the “Match between runs” option, in which the maximal retention time window was set to 0.7 min. Protein intensities were normalized using the in-built label-free quantification algorithm. Technical and biological replicates for each condition were defined as groups, label-free quantification intensity values were filtered for minimum value of 2 per group and transformed to log2 scale. Differences in protein levels between samples from unphosphorylated and phosphorylated columns were calculated as log2 fold change in protein intensity. Enriched proteins for each condition were determined by a log2 fold change ≤ 0.05 and ≥ 2.

### Immunofluorescent labeling

Cells and neurons were fixed with pre-warmed 4% PFA (#1040051000, Merck) PBS for 15 minutes, and rinsed with PBS. Cells were permeabilized and nonspecific binding blocked with PBS containing 0.5% TritonX100 (#T8655, US Biological) and 5% goat serum (#16210-072, Gibco) for 1 hour at room temperature. Primary antibodies (total volume 100 μl/coverslip; see Table 1) were incubated overnight at 4°C. The next day, coverslips were washed 2 x 10 minutes with PBS 0.1% Tween-20 followed by 2 x 10-minute washes with PBS. Secondary antibodies (total volume 100 μl/coverslip; see Table 1) were applied for one hour at room temperature, followed by 2 x 10-minute washes with PBS 0.1% Tween-20 and 2 x 10-minute washes with PBS. All antibodies were diluted in PBS containing 0.1% Tween-20 (#655205, Calbiochem) and 5% goat serum. Coverslips were mounted with Prolong Glass Antifade Mountant (#P36984, Invitrogen) and kept at 4°C.

### Analysis of PLPPR3 clusters in synapses

Images of DIV7 hippocampal neurons were acquired on Leica SP8 inverted confocal with 40x objective. Distal axons of healthy neurons with moderate expression of recombinant proteins were imaged. Channels were acquired sequentially, and acquisition filters were adjusted to avoid crosstalk of fluorophores. Z-stacks with a step size of 0.3 μm covering the entire axon were acquired.

The segmentation of PLPPR3 clusters and presynaptic clusters (defined by Synaptophysin-1 signal) was performed in an automated manner using a custom-written ImageJ macro (segmentation)^86^ and an additional Python script (distance measures), which can be found on GitHub (https://github.com/ngimber/axonal_cluster_workflow). Axons were segmented based on the Synaptophysin-1 signal as follows: uneven background was corrected by the ’Normalize Local Contrast’ function and images were smoothed (Gaussian blur, sigma = 280 nm) before binarization with the Triangle method.^87^ The ‘Closing’ operation was used (1.41 µm) to remove small gaps and linear filamentous structures were enhanced via ‘Ridge Detection’ before converting axons into skeletons with the ‘Skeletonize’ function. PLPPR3 clusters and the Synaptophysin-1 clusters (presynapses) were segmented with the following procedure: uneven background was compensated by ‘Rolling Ball Background Subtraction’ before applying a Gaussian Blur (sigma = 280 nm) for noise reduction and smoothing. The ‘Opening function’ was used to remove confocal shot noise and small objects. Clusters were binarized by applying an automated threshold (‘Moments’: Tsai’s method)^88^ and connected clusters were split into single clusters using the Watershed algorithm. Only clusters that overlapped with the segmented axons were used for further analysis. PLPPR3 clusters were categorized as ’within synapses’ if the cluster center was located within a radius of 0.7 µm (presynaptic diameter = 1.4µm)^89^ around the Synaptophysin-1 cluster center.

### Filopodia density measurements

N1E-115 cells were transfected with farnesylated GFP (independent membrane marker)^84^ combined with PLPPR3-Flag, PLPPR3-S351A-Flag or PLPPR3-S351D-Flag, and additionally labelled for F-actin (independent filopodia marker). Images of individual, non-overlapping cells were acquired on Leica SP8 inverted confocal with 63x objective. Channels were acquired sequentially, and acquisition filters were adjusted to avoid crosstalk of fluorophores. Z-stacks with a step size of 0.6 μm covering the entire cell were acquired. Filopodia density measurements were performed using an ImageJ macro developed by Joachim Fuchs.^30^ The macro automatically detects the borders of a cell and outlines it, providing a measurement of circumference. Filopodia are detected as intensity peaks along this outline. The analysis was carried out in the membrane marker channel. The ImageJ macro of this analysis is available at (https://github.com/jo-fuchs/Filopodia_Membrane_recruitment). Filopodia density experiment was replicated 3 times with different passages of cells.

### Prediction of phosphorylation sites and conservation analysis

Prediction of phosphorylation sites was carried out in NetPhos3.1 server (https://services.healthtech.dtu.dk/services/NetPhos-3.1/). The primary amino acid sequence of PLPPR3 intracellular domain (aa 284-716) was submitted, and phosphorylation of serine, threonine and tyrosine residues was selected for prediction. Only scores >.75 were displayed. To analyze conservation of phosphorylation sites across species, Constraint-Based Multiple Alignment Tool (COBALT, https://www.ncbi.nlm.nih.gov/tools/cobalt/cobalt.cgi?CMD=Web) was used. FASTA sequences of mouse (*Mus musculus*; Q7TPB0-1), human (*Homo sapiens*; Q6T4P5), zebrafish (*Danio rerio*; B0V0Y5), tropical clawed frog (*Xenopus tropicalis*; B1H174), rhesus monkey (*Macaca mulatta*; F7GER3) and chicken (*Gallus gallus*; A0A1D5PYJ6) from Uniprot database (https://www.uniprot.org/) were used. The sequences were aligned with default settings.

### Statistics and data visualization

Statistical analysis and data visualization was performed with Graphpad Prism 9.0.0 (121). Filopodia densities were compared with one-way ANOVA, changes in co-immunoprecipitation were compared with Wilcoxon test. Figures 1B, 2A, 2C, 4D and 5H were generated with Biorender.com.

## Acknowledgements

We thank Kerstin Schlawe and Kristin Lehmann for excellent technical assistance. We thank Katrin Büttner for help with cloning, Susanne Wegmann and Janine Hochmair (DZNE Berlin) for help with kinase assays, and Georg Braune for help with affinity chromatography. We thank Joachim Fuchs, Till Mack and Patricia Kreis (Charité Universitätsmedizin Berlin) for helpful discussions, and Alexandra Polyzou (University of Ioannina) for sharing her unpublished data with us. We are grateful to the Advanced Medical Bioimaging (AMBIO) Core Facility as well as to the NeuroCure Multi-user Microscopy Core Facility of Charité for assistance on microscopy and image analysis, and to Pico Caroni for sharing antibodies with us. Parts of this manuscript were published as a monograph to obtain the doctoral degree (available at https://refubium.fu-berlin.de/handle/fub188/41517). This work was funded by DFG under the Collaborative Research Centre SFB958 project A16 and Z02.

