## Supplemental Figures for "Phosphorylation of PLPPR3 membrane proteins as signaling integrator at neuronal synapses"

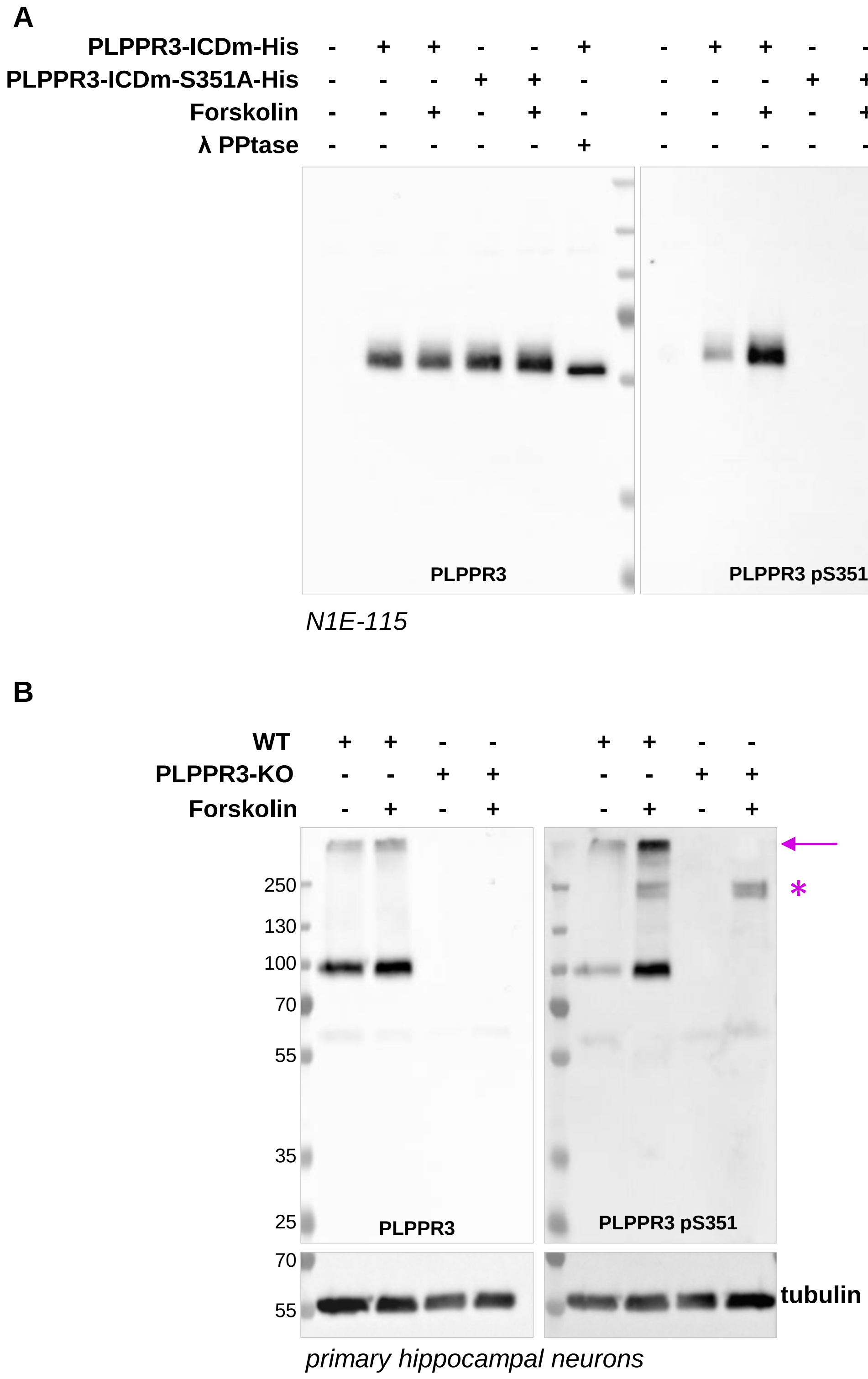


*Figure S1.* Validation of the PLPPR3 pS351 antibody. (A) Specificity of the PLPPR3 pS351 antibody towards recombinant PLPPR3 ICDm. PLPPR3 ICDm variants were expressed in N1E-115 cells, proteins were extracted, and the lysates were analyzed by western blot. Forskolin stimulation (30 µM, 5 minutes) was performed before lysis. Λ phosphatase treatment was performed at 30°C for 30 minutes on phosphatase inhibitor-free samples after cell lysis. (B) Specificity of the PLPPR3 pS351 antibody towards endogenous PLPPR3. Primary hippocampal neurons were lysed at DIV9, proteins were extracted, and the lysates were analyzed by western blot. Forskolin stimulation (30 µM, 5 minutes) was performed before lysis. Unspecific bands are annotated with an asterisk. Higher-order oligomerized forms of PLPPR3 are annotated with an arrow.

| Gene name | Cleavage coverage [%] | Nr of unique peptides | Log2 intensity deP | Log2 intensity P | Log2 fold change P vs deP | Species |
|---|---|---|---|---|---|---|
| Basp1 | 96 | 25 | 22 | 29 | 7 | mouse |
| Atp5b | 82 | 26 | 25 | 30 | 5 | mouse |
| Hist1h1e | 40 | 3 | 27.22 | 26.54 | -0.8 | mouse |

*Figure S2.* List of potential interaction partners from the PLPPR3 pS351 peptide affinity chromatography mass spectrometry analysis. deP = non-phospho peptide, P = phospho-peptide column.
